# Caecilians maintain a functional long-wavelength-sensitive cone opsin gene despite signatures of relaxed selection and more than 200 million years of fossoriality

**DOI:** 10.1101/2025.02.07.636964

**Authors:** María José Navarrete Méndez, Sina S. Amini, Juan Carlos Santos, Jacob Saal, Marvalee H. Wake, Santiago R. Ron, Rebecca D. Tarvin

## Abstract

Vision is tuned to animals’ ecologies, evolving in response to specific light environments and visual needs. Transitions to fossorial lifestyles impose strong selective pressures favoring adaptations for underground life, such as increased skull ossification and reduced eye protrusion. Fossoriality may simultaneously relax constraints on vision leading to diminished visual capabilities. Caecilians (Gymnophiona)—specialized, fossorial amphibians—possess reduced eyes covered by skin or bone. For years, these traits, along with the presence of a single photoreceptor expressing one functional opsin gene, have been interpreted as evidence of limited vision, including an inability to focus or perceive color. Our results challenge these assumptions: we identified the long-wavelength-sensitive (*LWS*) opsin gene in 13 species of caecilians spanning 8 of 10 recognized families. Molecular evidence indicates that *LWS* is intact and transcribed in the eye of at least one species (*Caecilia orientalis*). However, the specific photoreceptor type expressing *LWS* remains uncertain, as our survey of cone phototransduction genes revealed a mosaic of losses, and anatomical observations from five families did not conclusively identify cone-like cells, though they revealed highly organized retinae even in families with vestigial eyes. Altogether, our results suggest that vision in caecilians may be underestimated and the role of light perception in their ecology is possible.

**TEASER TEXT:** The colonization of low-light environments exerts selective pressure to enhance non-visual senses while relaxing selection on vision, often leading to opsin gene loss. We present compelling evidence for the retention of a cone opsin gene (*LWS*) in caecilians—a fossorial amphibian lineage with a 200-million-year history of burrowing and reduced eyes. Despite signs of relaxed selection, *LWS* appears functional, echoing patterns in other low-light-adapted vertebrates. Additionally, the loss of key cone phototransduction genes and absence of clear cone cells raises the possibility of *LWS* expression in rods or a functional change in its use. Our study provides an evolutionary framework for investigating *LWS* retention in fossorial vertebrates.

## 1. INTRODUCTION

Strong selection for adaptation to new habits, such as fossoriality (i.e., organisms that live underground), results both in gain and loss of traits (Nevo, 1979). For example, some burrowing frogs evolved spades through the enlargement of metatarsal tubercles (Emerson, 1976), yet their altered body and limbs trade off with jumping ability (Vidal García et al., 2014). The star-nosed mole (*Condylura cristata*) evolved a novel mechanosensory structure that allows them to “see” prey (Catania, 1999; Holmberg, 2022) despite having small, underdeveloped eyes (Carmona et al., 2010). In these and other animals, subterranean life intensifies selection pressures for adaptations that facilitate life underground and relaxes selection for traits that are no longer as relevant.

Although relaxed selection can lead to trait degeneration, it is rare that traits are ever completely lost because they may be developmentally constrained, still possess some functional value, or cost little to maintain (Sadier et al., 2022). Partial eye degeneration without complete loss is a common characteristic of fossorial, deep-sea, and cave-dwelling vertebrates. For example, the European mole (*Talpa europaea*) and Iberian mole (*Talpa occidentalis*) have organized retinae with rod and cone cells despite being fully subterranean (Carmona et al., 2009; Glösmann et al., 2008); *T. occidentalis* never even opens its eyelids. Eyeless populations of the cave-dwelling fish *Astyanax mexicanus* develop a layered retina during their embryonic stage, retaining the ability to exhibit light-entrained retinomotor rhythms even as their eyes undergo extensive ontological degeneration (Emam et al., 2020; Espinasa & Jeffery, 2006).

Eye degeneration can be coupled at the molecular level with the loss of visual opsin genes that encode sensitive photoreceptor proteins (Bowmaker, 2008; Emerling & Springer, 2014; Lin et al., 2024). For example, the Cape golden mole (*Chrysochloris asiatica*), which is almost exclusively subterranean and has subcutaneous eyes, exhibits one of the highest numbers of pseudogenes among fossorial mammals, including losses of cone opsins and several other genes involved in the phototransduction cascade (Emerling & Springer, 2014). However, opsin genes may be lost even when ocular anatomy remains well-developed, including features such as an organized and layered retina (Foureaux et al., 2010; Wake, 1985; Warrant, 2000). Across vertebrates, independent losses of opsins have been proposed as adaptations to specific visual demands shaped by behavioral and ecological shifts (W. L. Davies, 2011). For instance, at least 10% of mammalian species are thought to lack functional copies of the short-wavelength sensitive opsin gene (*SWS1*), and all placental mammals have lost the middle-wavelength sensitive opsin gene (*RH2*)–patterns attributed to nocturnal lifestyles and/or ancestry (W. I. L. Davies et al., 2012; Hunt et al., 2009; Jacobs, 2013).

Interestingly, although opsins have been lost multiple times in vertebrates, others tend to persist (W. L. Davies, 2011), even when environmental conditions seem unsuitable for their spectral sensitivity. Among the five visual opsin genes found in vertebrates (i.e. *RH1*, *RH2*, *LWS*, *SWS1*, *SWS2*), the *LWS* opsin exhibits peak sensitivity to the longest wavelengths within the visible spectrum (560 – ∼570 nm, corresponding to red-green hues, though sensitivity can extend to wavelengths beyond the peak absorbance; (Liebman & Entine, 1968; Margetts et al., 2024) and is one of the most consistently retained across vertebrates, including species adapted to low-light environments (W. I. L. Davies et al., 2012; Ford et al., 2024; Musilova et al., 2021).

Ancestors of caecilians, a group of limbless amphibians, began their transition to a subterranean lifestyle more than 250 million years ago (MYA) through elongation of the trunk, fusion of cranial bones, and reduction of limbs (Jenkins & Walsh, 1993; Santos et al., 2020). In extant caecilians (with most recent common ancestor 190 MYA; Kligman et al., 2023) all species lack limbs, have eye sockets that are covered by skin or by skin and bone, and lack the eye muscles for accommodation (Wake, 1985). Visual system reduction correlates broadly with fossorial specialization (Himstedt, 1995; Kamei, San Mauro, et al., 2012): early-diverging, less fossorial caecilian lineages like ichthyophiids and rhinatrematids retain better-developed eyes, while taxa with more derived fossorial adaptations–such as *Boulengerula boulengeri*–often exhibit extreme ocular reduction (Mohun & Wilkinson, 2015; Wake, 1985). However, certain aspects of visual system anatomy—such as the degree of degeneration in extrinsic eye muscles, lens structure, and even retinal organizations—display a mosaic pattern of trait loss across species, independent of phylogeny, ecology, or cranial ossification (Mohun & Wilkinson, 2015; Wake, 1985).

In spite of these morphological constraints, evidence from anatomy, physiology, and molecular biology has shown that visual systems in caecilians have remained somewhat functional (Dünker, 1998; Mohun et al., 2010; Mohun & Wilkinson, 2015; Wake, 1985). For example, negative phototactic behavior has been reported in *Ichthyophis cf. kohtaoensis* (Himstedt, 1995; Mohun et al., 2010). Lens morphology appears to be adapted to environment, as the aquatic/semi-aquatic family Typhlonectidae possesses spherical lenses (similar to most aquatic vertebrates), in contrast to the ellipsoid lenses typical of most terrestrial caecilians (Wake, 1985). Furthermore, the rod phototransduction pathway used for scotopic vision remains functional in Gymnophiona, as evidenced by electroretinography and microspectrophotometry analyses that detected a visual pigment with peak absorbance near 500 nm (consistent with the spectral properties of *RH1*) in *Ichthyophis* cf. *kohtaoensis* (Himstedt, 1995; Mohun et al., 2010)*, Rhinatrema bivittatum, Geotrypetes seraphini,* and *Typhlonectes natans* (Himstedt, 1995; Mohun et al., 2010). The identity of this pigment was later confirmed in two of these species (i.e. *I.* cf. *kohtaoensis* and *T. natans*) as the rod opsin *RH1* (Himstedt, 1995; Mohun et al., 2010).

Moreover, although caecilian retinas are thought to lack cone cells (Mohun & Wilkinson, 2015; Wake, 1985), a gene encoding the long-wavelength-sensitive cone opsin (*LWS*) was recently identified in three published caecilian genomes from the families Rhinatrematidae, Dermophiidae, Siphonopidae (Lin et al., 2024).

Here we investigated the evolution of the two visual opsin genes known to be present (*LWS* and *RH1*) in caecilians and explore why *LWS* has been maintained without loss or pseudogenization for hundreds of millions of years despite significant eye degeneration and an ecological context wherein it would otherwise appear superfluous. Given that intact copies of the *LWS* gene have been identified in three distantly related caecilian species exhibiting varying degrees of skull ossification and eye reduction (Lin et al., 2024), we hypothesize that *LWS* was present in the common ancestor of caecilians and has been maintained in all extant species, independent of eye exposure or anatomical complexity. To test this, we reconstructed the evolution of eye musculature and exposure using ancestral state reconstruction to infer the ancestral condition, trace the trajectory of eye degeneration, and evaluate the persistence of *LWS* in this anatomical context.

Additionally, given that caecilian *LWS* genes exhibit intact open reading frames without premature stop codons, we hypothesize that they remain functional and may be expressed in the eye, with absorbance properties resembling those of long-wavelength-sensitive opsins in amphibian cones. To test this, we sequenced the eye transcriptome of one caecilian species and used a machine learning approach to estimate the absorbance peak of caecilian *LWS* based on its amino acid sequence.

Finally, to investigate whether cone photoreceptors capable of expressing *LWS* persist in caecilians, we used two complementary approaches. First, we examined the genomes and transcriptomes of caecilian species for genes associated with the cone signaling cascade, under the hypothesis that if *LWS* is expressed in cones, the downstream components of the cone phototransduction pathway should also be present. Second, we reviewed histological material to search for evidence of cone photoreceptors. We further hypothesize that selection analyses will reveal evolutionary constraints on *LWS* that parallel patterns observed for the rod opsin gene *RH1*, thereby supporting its continued physiological relevance. As one of the earliest vertebrate lineages to adopt a subterranean lifestyle, caecilians provide a unique opportunity to explore the evolution and persistence of light perception in fossorial environments.

## 2. METHODS

### 2.1 ANCESTRAL CHARACTER STATE RECONSTRUCTION

We estimated ancestral character states for eye musculature and exposure using character data mainly from Wake (1993), but also, for a few species, from Norries & Hughes (1918), Taylor (1968, 1969), and Maciel and Hoogmoed (2011) (see Supplementary Table 1). The reconstructions were performed using stochastic character mapping in *phytools* v2.0 (Revell, 2024) on a time-tree obtained from the San Mauro et al. (2014) matrix of mitochondrial genomes. The time-tree was inferred using BEAST v2.7.8 (Bouckaert et al., 2019) under a strict clock and the model of DNA evolution HKY. See Supplementary Methods section 2.1 for more details.

### 2.2 MORPHOLOGY AND SEARCH FOR CONE PHOTORECEPTORS

#### 2.2.1 Histology

The examined sections are archival material accessioned in the MVZ that were originally prepared by MHW in the 1980s. Details on sample preparation, staining, and storage are provided in Supplementary Methods section 2.2.1. Sampling was opportunistic, representing all species with available histological sections suitable for visualizing eye morphology. More information about these species and accession information can be found in Supplementary Table 2 (see also (Wake, 1985).

#### 2.2.2 Photomicrography

Eyes were observed with a Laxco microscope, where we examined sections with intact photoreceptors and retina. Photographs were taken at standard low, medium, and high magnifications using a Laxco SeBaCam5C digital microscope camera, and rendered using SeBaView software v3.7 (Laxco, Inc.). A stage micrometer was used for measurements and to calibrate scale bars.

#### 2.2.3 Search for cones and description of the retina

We visually surveyed all images for photoreceptor cells with morphology characteristic of cones (i.e., bulbous ellipsoids, tapered outer segments that are much thinner than their inner segments) or otherwise distinct from the rods that predominate the retina. Slides of *Epicrionops* sp., *Typhlonectes compressicauda*, *Ichthyophis kohtaoensis*, *Dermophis mexicanus*, and *Scolecomorphus kirkii* specimens were selected to highlight taxonomic and morphological diversity of retinae across Gymnophiona due to their taxonomic disparity and well-preserved morphology. Standard descriptions of the retinae were made following (Mohun & Wilkinson, 2015; Wake, 1985) and (Himstedt, 1995; Mohun et al., 2010). Details on measurement methodology are provided in Supplementary Methods section 2.2.3.

### 2.3 FURTHER VALIDATION OF A CAECILIAN *LWS* AND ASSESSMENT OF ITS POTENTIAL FUNCTIONALITY

#### 2.3.1 Generating an eye transcriptome

We collected one individual of *Caecilia orientalis* in 2020 at Yanayacu Biological Station in Napo, Ecuador (S 0.5984097, W 77.8907409, 2143 m.a.s.l.) under permit MAE-DNB-CM-2015-0025-M-0001 issued by the Ministerio del Ambiente of Ecuador to Universidad Católica del Ecuador.

We euthanized the specimen using commercial Roxicaine anesthetic (lidocaine) and immediately dissected both eyes which were preserved in RNAlater. The specimen is deposited in the collection of Museo de Zoología of the Pontificia Universidad Católica del Ecuador (QCAZ), with specimen number QCAZ:A:77338. Tissue samples were sent to Macrogen Inc. (Seoul, South Korea) for RNA extraction, library preparation using the Illumina TruSeq Stranded mRNA LT Sample Preparation Kit, and sequencing on the Illumina NovaSeq6000 to 100 million paired-end 100-bp reads.

Read quality was assessed using FastQC (Andrews, 2010). We used TrimGalore! (https://www.bioinformatics.babraham.ac.uk/projects/trim_galore/) to remove adapters, low-quality bases, and short reads. Transcriptome assembly was then conducted *de novo* using Trinity v2.13.2 (Grabherr et al., 2011), incorporating all paired-end reads with default settings. The resulting assembly was filtered using SeqClean (Chen et al., 2007). To annotate protein-coding genes, TransDecoder v2.026 (Haas, n.d.) was used with default parameters and homology to UniProt was then used to infer functions of predicted genes with NCBI-BLASTP v2.2.30+ (E-value = 1e−6) and Interproscan 5.35 (Jones et al., 2014) to detect proteins with known functional domains. Finally, we used Salmon v1.10.3 (Patro et al., 2017) to quantify transcript abundances of opsin genes. First, we built a Salmon index for our transcriptome using the index command with default parameters for k-mer length. Next, we quantified the paired-end reads against the previously generated index using the quant command. The output gene counts were summarized as normalized transcript abundances using DESeq2 v3.11 (Love et al., 2014).

#### 2.3.2 PCR of LWS in additional caecilian species

To validate whether *LWS* existed in a broader phylogenetic and ecological sampling of caecilians, we obtained tissue samples from terrestrial and aquatic species exhibiting varying degrees of fossoriality and reports of above- ground behavior. We included samples of species from early branching lineages that exhibit lesser degrees of adaptation to fossoriality including *Epicrionops bicolor*, *E. petersi* (Rhinatrematidae), as well as more phenotypically derived species such as *Caecilia orientalis* (Caecilidae) and *Microcaecilia albiceps* (Siphonopidae) from specimens at QCAZ. Additional tissues were obtained from the Museum of Vertebrate Zoology (MVZ) at the University of California, Berkeley, including terrestrial species that inhabit moist, loose soils with varying levels of eye coverage such as *Dermophis mexicanus*, *Geotrypetes seraphini*, *Gymnopis multiplicata* (Dermophiidae), as well as *Ichthyophis kohtaoensis* (Ichthyophiidae) which has aquatic larvae and terrestrial adults, and the fully aquatic *Typhlonectes natans* (Typhlonectidae). Tissues from species with markedly reduced eyes, such as *Boulengerula boulengeri* (Herpelidae) and *Scolecomorphus vittatus* (Scolecomorphidae), were kindly loaned from the California Academy of Sciences. We attempted to obtain tissues that match our morphological material to the extent possible (at least to genus). Together, these samples represent the full range of diversity in caecilian eye morphology. See Supplementary Table 3 for the accession numbers.

DNA extraction, amplification, and sequencing was carried out at PUCE for QCAZ specimens, and at the Evolutionary Genomics Lab (EGL), UC Berkeley, for MVZ and CAS specimens. Fragments of *LWS* were amplified using new primer sets designed from an alignment of amphibian opsins. The new LWS_542R (5’-GCA GAC CAA ACC CAG GAG AA-3’) and LWS_117F (5’ TTT TGA GGG MCC CAA CTA CC -3’) primers amplify a region that includes part of exon I, the intron between exon I and II, and part of exon II. Specifics about the extraction and amplification protocols are provided in the Supplementary Material Methods section 2.3.2. Additionally, an evaluation of primers used in previous attempts that did not yield successful amplification of *LWS* in caecilians (Mohun et al., 2010) is provided in Supplementary Material Methods section 2.4.

PCR products from QCAZ samples were sequenced at Macrogen Inc. (Seoul, South Korea); MVZ and CAS samples were sequenced at the UC Berkeley DNA Sequencing Facility. We assembled forward and reverse sequences into consensus sequences using default parameters in Geneious Prime 2022.1.1 and then aligned them against each other using MUSCLE (Edgar, 2004). To confirm the identity and quality of the new *LWS* sequences, we inferred gene trees for *LWS* and *RH1* in IQTREE2. Sequences were partitioned by coding and non-coding regions and also by codon position. The best-partition model was estimated using the -TESTMERGE option (Chernomor et al., 2016; Kalyaanamoorthy et al., 2017). Bootstrap values were calculated using 500 replicates (-b 500) and were plotted on the best likelihood tree. See Supplementary Data 1 for input and output files (Supplementary Data 1-7 are archived at Dryad—DOI: 10.5061/dryad.h18931zxf).

#### 2.3.3 Estimation of the wavelength of maximum absorbance (***λ***_max_) of *LWS* and *RH1*

Given that the *LWS* copies identified in caecilian genomes contain open reading frames without stop codons (i.e., considereded intact), we hypothesize that the gene is functional and that the protein encoded by *LWS* will exhibit absorbance spectra within the expected ranges (560–570 nm; (W. I. L. Davies et al., 2012). To test this hypothesis, we used a machine-learning model (Frazer et al., 2024) to estimate the wavelength of maximum absorbance (λ_max_) for five full-length *LWS* sequences from the caecilian species included in our study incorporating data available on GenBank (*Rhinatrema bivittatum*, XM_029609805.1; *Microcaecilia unicolor,* XM_030192059.1; *Geotrypetes seraphini*, XM_033922369.1; *Ichthyophis bannanicus*, GCA_033557465.1 hereafter *I. kohtaoensis* following Nishikawa et al., 2021) and our *C. orientalis* eye transcriptome. Additionally, we estimated absorbance values for the *RH1* sequences of the same species listed above to validate the accuracy of the machine-learning model, since experimental absorbance estimates for this opsin have been reported for *G. seraphini* and *R. bivittatum* (Mohun et al., 2010). We expected the estimated λ_max_ values for *RH1* to fall within the typical range for this opsin (∼500 nm; W. L. Davies, 2011) and to closely approximate experimental MSP data. Furthermore, we increased our phylogenetic sampling by including *LWS* and *RH1* sequences from other relevant amphibian and vertebrate species. For this purpose, we explored three types of sequence archives: NCBI sequence references, unannotated transcriptomes, and published genomes. The methods used to retrieve *LWS* from each of these datasets are described in more detail in the Supplementary Material section 2.3.3 Our final dataset included 19 species of Anura and 8 species of Caudata (see Supplementary Table 3 for details). The nucleotide matrices were manually inspected to remove any ambiguities and the sequences were translated to amino acids for downstream analysis (see Supplementary Data 2).

Point estimations of λ_max_ for *LWS* and *RH1* were predicted using the vertebrate model trained with the VPOD_vert_het_1.0 dataset available in the Visual Physiology Opsin Database (https://github.com/VisualPhysiologyDB/visual-physiology-opsin-db). This model demonstrated the best predictive ability, with the highest 10-fold cross-validation R² (0.968) and the lowest mean absolute error (6.56 nm) among all the models compared in the machine learning analysis (Frazer et al., 2024). The training dataset included 721 vertebrate opsins (UVS, SWS, MWS, LWS, RH1, RH2) spanning wild-type, mutant, and chimeric sequences with λ_max_ values ranging from 350 to 611 nm (Frazer et al., 2024).

Additionally, a literature search was conducted to identify available microspectrophotometry (MSP) data for the species included in our machine-learning analysis. In some instances, when values were not available for a particular species, we used data from a closely related species within the same genus (Supplementary Table 4). We then performed a direct comparison between the MSP data and the machine-learning estimations using a linear correlation analysis (Pearson’s correlation) using the *lm* and *cor* test functions in Rstudio v2024.9.1.394 and calculated the mean absolute error to quantify the discrepancy between the absorbance values obtained from the two methodologies.

#### 2.3.4 Search for presence and expression of rod and cone phototransduction cascade genes

The presence of *LWS* in caecilians contrasts with the lack of morphological evidence for cone photoreceptors, which are the typical cellular structure for *LWS* expression. We hypothesized that if *LWS* is expressed and functional in cones, intact cone phototransduction genes should also be present and expressed, and thus detectable in both the genomes and transcriptomes of caecilians. Alternatively, if caecilian retinas contain only rod photoreceptors, as previously reported, and *LWS* is potentially co-expressed with *RH1* in rods, we would expect to find only rod-associated phototransduction genes, with cone-specific genes either pseudogenized, absent, or not expressed in the eye transcriptome. To test this, we searched for rod and cone phototransduction genes in NCBI reference databases and publicly available genomes of three anuran species: *Xenopus tropicalis* (GCF_901001135.1), *Nanorana parkeri* (GCF_000935625.1), and *Pyxicephalus adspersus* (GCA_032062135.1); two salamanders: *Ambystoma mexicanus* (GCA_040938575.1) and *Pleurodeles waltl* (GCA_031143425.1); and three caecilians: *Microcaecilia unicolor* (GCF_901765095.1), *Rhinatrema bivittatum* (GCF_901001135.1), and *Geotrypetes seraphini* (GCF_902459505.1). Additionally, we searched for these genes in our newly generated eye transcriptome from *Caecilia orientalis* (PRJNA1181370) (Supplementary Table 3). The canonical visual phototransduction cascade in vertebrates involves three main components: (1) the heterotrimeric G-protein transducins (in rods: *GNAT1*, *GNB1*, *GNGT1*; in cones: *GNAT2*, *GNB3*, *GNGT2*); (2) the phosphodiesterase enzymes (in rods: *PDE6A*, *PDE6B*, *PDE6G*; in cones: *PDE6C*, *PDE6H*), and (3) the cyclic nucleotide-gated channels (in rods: *CNGA1*, *CNGB1*; in cones: *CNGA3*, *CNGB3*) (Lamb, 2013).

The identified sequences of our genes of interest were downloaded and used as reference queries to search our *de novo*–assembled transcriptome data, which had been formatted as a custom BLAST database using the default parameters for *makeblastdb* (v2.16.0+). All the obtained sequences were translated into amino acids and aligned using MAFFT v7.490 (Katoh & Standley, 2013), and the alignments were manually inspected in Geneious Prime 2022.1.1 (https://www.geneious.com) for the presence of premature stop codons. Finally, following the methodology described in section 2.2.1, we used the previously generated Salmon index of the eye transcriptome to quantify transcript abundances of visual phototransduction genes and assess whether their expression levels mirror those of the opsin genes. When multiple isoforms were reported for a given gene, we summed the expression levels of all isoforms. Although some isoforms may result from assembly artifacts, we opted to include all of them to avoid making an arbitrary choice about which isoform to retain.

### 2.4 SELECTION ANALYSES

The presence of intact *LWS* genes without nonsense and frameshift mutations in five phylogenetically diverse caecilian species supports the idea that *LWS* is an ancestrally retained feature despite a long fossorial history. Thus, we hypothesized that *LWS* is under evolutionary constraint, similar to *RH1*, indicating a physiological role similar to its ancestral function. To test this, we conducted a series of selection analysis on *LWS* and *RH1* in Gymnophiona and other amphibian species using the sequence data described in section 2.2.3, including transcriptomic data from *C. orientalis* and *LWS* fragments obtained via Sanger sequencing. In total, our sampling included 13 caecilian species spanning eight families (Supplementary Table 3). An updated and unrooted version of the phylogenetic tree presented by (Jetz & Pyron, 2018) served as a backbone for conducting selection analysis in the codeml program of PAML v4 (Yang, 2007).

Sequences for *LWS* and *RH1* were aligned using MAFFT (Katoh & Standley, 2013) followed by manual editing to remove terminal stop codons in Mesquite v3.6 (Maddison & Maddison, 2015). For *LWS*, we ran analyses on two types of datasets, a complete dataset including the full coding sequence of the gene with smaller sampling for Gymnophiona (368 aa; Anura = 19 spp, Caudata = 8 spp, Gymnophiona = 5 spp) and an incomplete dataset with more species for Gymnophiona but fewer sites (132 aa; Anura = 19 spp; Caudata = 8 spp, Gymnophiona = 13 spp). For *RH1*, we only ran analyses using the complete sequences (354 aa; Anura = 18 spp, Caudata = 6 spp, Gymnophiona = 5 spp).

We initially conducted a broad phylogeny-wide analysis to assess the strength of purifying selection on *LWS* and *RH1*. We used the M0 model, which assumes a single ω (dN/dS) ratio across all sites and lineages. To represent the null hypothesis of neutral evolution, we fixed ω at 1 (fix_omega = 1; omega = 1), thereby constraining the model to assume no selection. To test our hypothesis that *LWS* has been maintained under purifying selection, we ran an alternative model in which ω was estimated from the data (fix_omega = 0; omega = 0.5). We then compared the likelihoods of the two models using a likelihood ratio test (LRT).

We also inferred selection patterns and tested for site-specific positive selection for each gene comparing the support for different models (M0, M1a, M2a, M2a_rel, M3, M7, M8a, and M8). We used the Bayes Empirical Bayes (BEB) approach to identify individual sites with a posterior probability >90% of being in the positively selected class of sites. Additionally, we compared the fit of clade models C and D (CmC vs. CmD) (Bielawski & Yang, 2004) to M3 and M2a_rel to investigate long-term shifts in selection pressures associated with the evolution of fossoriality in caecilians (Weadick & Chang, 2012). Both clade models assume three site classes, with the third class modeling divergent selection among clades; however, unlike CmC, CmD does not constrain the ω estimates for any site class. The best model was identified using likelihood ratio tests (LRTs) with a χ^2^ distribution.

Additionally, we conducted a series of analyses in HYPHY, implemented on the Datamonkey web server (Delport et al., 2010). FEL (Fixed Effect Likelihood; Kosakovsky Pond & Frost, 2005) and FUBAR (Fast, Unconstrained Bayesian AppRoximation; Murrell et al., 2013) were employed to estimate positive or purifying selection across sites by comparing single estimates of dN and dS values for each site. FUBAR uses a hierarchical Bayesian method rather than maximum likelihood, making it more sensitive than FEL to weaker signatures of positive selection (i.e., when ω is close to 1) (Murrell et al., 2013). We used BUSTED (Branch-site Unrestricted Statistical Test for Episodic Diversification; Murrell et al., 2013) to identify episodic positive selection at a fraction of sites. BUSTED allows for branch-to-branch variation across the entire phylogeny. Finally, we ran RELAX (Wertheim et al., 2015) to detect relaxation of selective constraint. This methodology requires an *a priori* specification (e.g., partition) of a subset of branches into foreground and background. In our case, the foreground comprised the crown Gymnophiona and its stem branch, while anurans and salamanders were designated as the background. RELAX estimates a selection intensity parameter (*k*), where *k* < 1 indicates relaxed selection and *k* > 1 indicates intensified selection, based on shifts in ω values between test and reference lineages.

We report results from the selection analyses using amino acid site numbers based on bovine rhodopsin (NP_001014890.1; alignments with bovine rhodopsin are available in the Supplementary Data 3). Selection results are reported by referring to the location of each amino acid according to the 3D structure of opsins, which encompasses seven transmembrane domains (TMD I–VII), three extracellular domains (ECD I–III), and the amino- and carboxyl-termini (N and C) (Palczewski et al., 2000). All input and output files for PAML and HYPHY selection analyses are available in Supplementary Data 4.

Finally, we examined *LWS* amino acid alignments at key spectral tuning sites to characterize the residue identity and variation at positions previously shown to influence the absorption properties of *LWS* and *RH1* opsins in vertebrates (Asenjo et al., 1994; Yokoyama, 2002a; Yokoyama & Radlwimmer, 2001a). We then compared the amino acid variants at these sites with those reported for amphibians in the recent comprehensive analysis by (Schott et al., 2024).

## 3. RESULTS

### 3.1 ANCESTRAL CHARACTER STATE RECONSTRUCTION

Ancestral character reconstruction of eye musculature shows that the ancestral condition for caecilians was the presence of all six eye muscles (Figure 1A). The most highly supported scenario suggests a retention of all muscles across most lineages, except for at least six independent losses of muscles. Notably, there was a loss from six to zero muscles, independently, in the MRCA of *Boulengerula* and *Gymnophis* The preferred model was equal rates (AIC 69.60), followed by ordered (AIC 80.46), and all rates different (AIC 114.98). The transition rate for the equal rates model was 0.0012.

**FIGURE 1.**
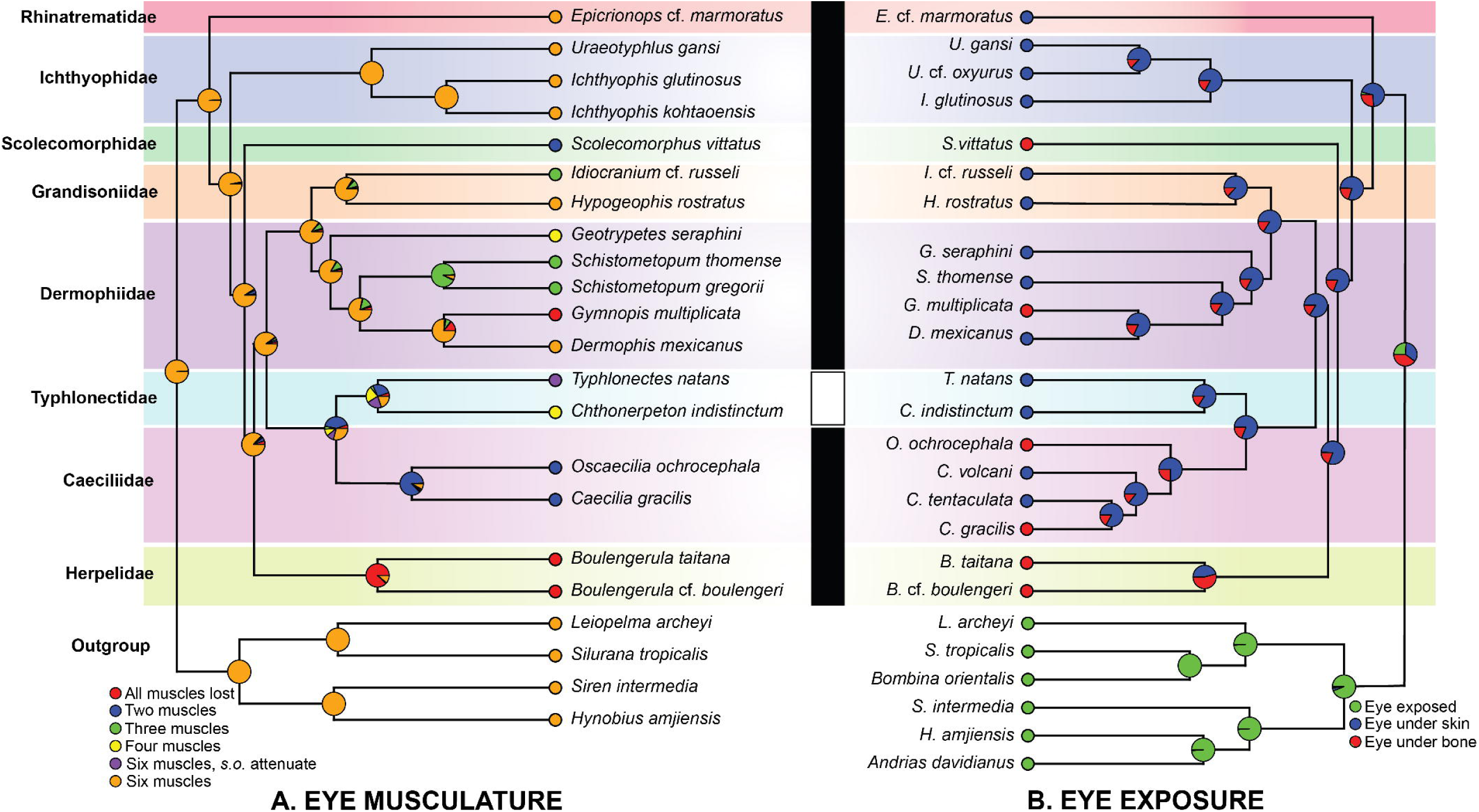
Posterior probabilities at each ancestral node for eye exposure and eye musculature condition. Probabilities were obtained from stochastic character mapping using model weights to sample three (A) or two (B) different Mk trait evolution models. The vertical bar indicates habitat type: black for terrestrial burrowers, white for aquatic and semi-aquatic burrowers. See Supplementary Methods section 2.1 for details.

Reconstruction of eye exposure shows that the ancestral condition for Gymnophiona is eye covered by skin (Figure 1B). From that ancestral condition, there have been five independent gains of eye covered by bone in *Boulengerula*, *Caecilia gracilis*, *Gymnopis*, *Oscaecilia* and *Scolecomorphus*. Note that the character is labile in the caecilids *C. gracilis* and *O. ochrocephala* (See Supplementary Material section 4.3). The preferred model was equal rates (AIC 46.09), followed by all rates different (AIC 46.37). The transition rate for the equal rates model was 0.00171.

### 3.2 CAECILIAN RETINAL MORPHOLOGY

282 photomicrographs of retinae representing 8 of the 10 families of Gymnophiona were examined (see Supplementary Table 2 for specimen information and Supplementary Data 5 for all photomicrographs). A subset of images with clear, well preserved morphological characters were selected to highlight retinal diversity across Gymnophiona as well as the finding of scattered cone-like photoreceptors. (Figures 2 and 3). All specimens examined had organized retinae containing all the cellular layers typical for visual vertebrates except *Boulengerula boulengeri* and *Scolecomorphus uluguruensis,* but this may be a result of poor preservation, particularly in *S. uluguruensis*. Species of the primarily aquatic family Typhlonectidae (*Chthonerpeton indistinctum, Typhlonecte*s *compressicauda*) possess spherical lenses, while terrestrial species exhibit ellipsoid lenses (e.g., *Epicrionops* sp*., Dermophis mexicanus)* except in *Caecilia occidentalis* and all species which have the orbit covered by bone (*Scolecomorphus kirkii, S. uluguruensis, Boulengerula boulengeri, Gymnopis multiplicata)*, where the lens is amorphous. Quantitative measures of the retina varied considerably among these taxa (Table 1). We found cone-like cells in 4 of 14 species (*Epicrionops* sp*., Dermophis mexicanus, Ichthyophis kohtaoensis*, *Scolecomorphus kirkii)*, but note that the sections for most specimens did not preserve the photoreceptor layer sufficiently to allow observation of individual photoreceptor morphology (See Supplementary Data 5). Detailed descriptions of eye morphology for the evaluated species are presented in Supplementary Material section 3.2.

**FIGURE 2.**
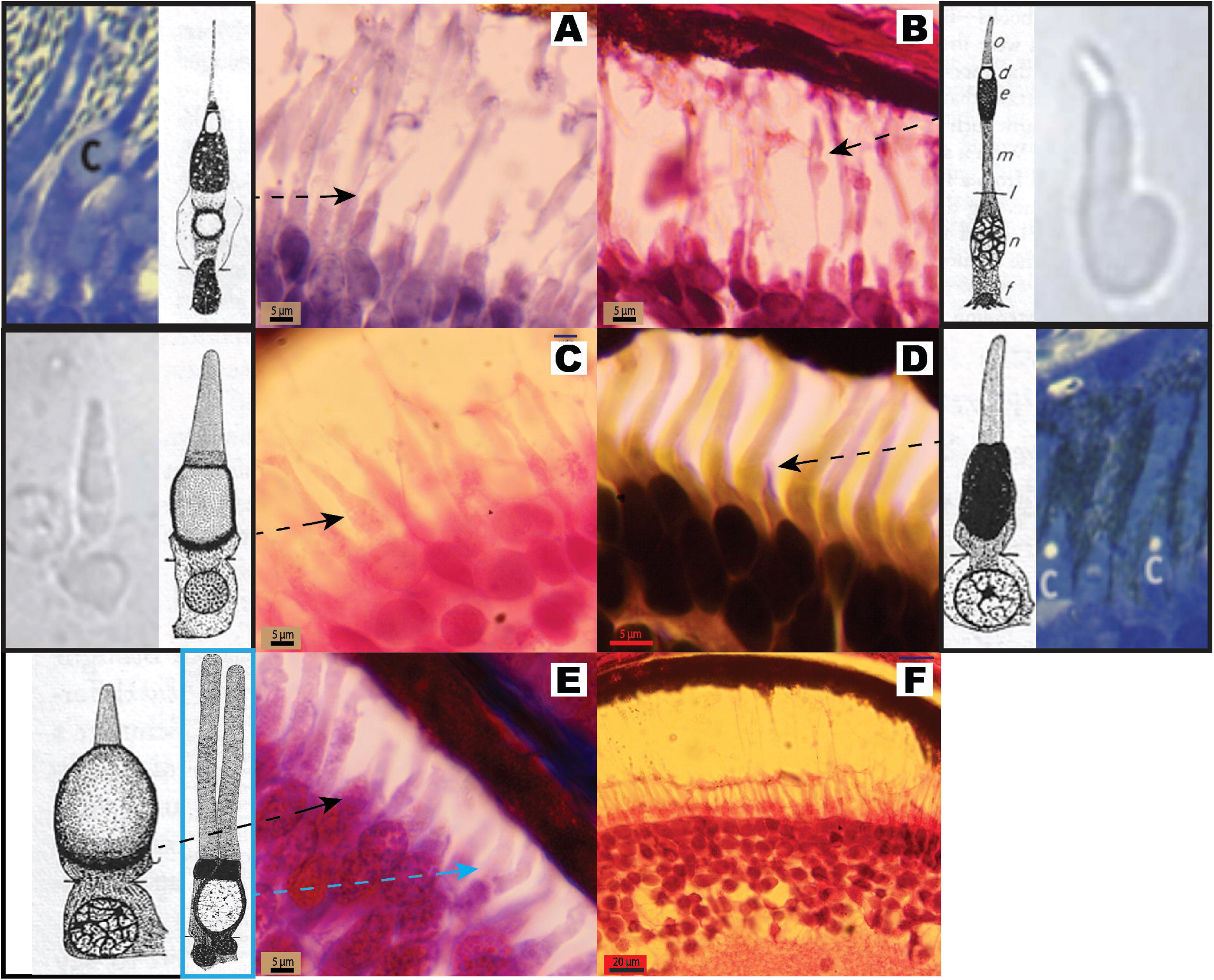
Photomicrographs of retinae from five species of caecilians at low and medium magnification. (A, B) *Epicrionops sp.* (C, D) *Ichthyophis kohtaoensis* (E, F) *Dermophis mexicanus* (G, H) *Typhlonectes compressicauda* (I, J) *Scolecomorphus kirkii.* Scale bars are 100 µm (A, C, E, G, I) and 20 µm (B, D, F, H, J). Letter abbreviations in (B): GCL - Ganglion cell layer; IPL - Inner Plexiform Layer; INL - Inner Nuclear Layer; OPL - Outer Plexiform Layer; ONL - Outer Nuclear Layer; IS - Inner Rod Segment; OS - Outer Rod Segment; PE - Pigment Epithelium.

**FIGURE 3.**
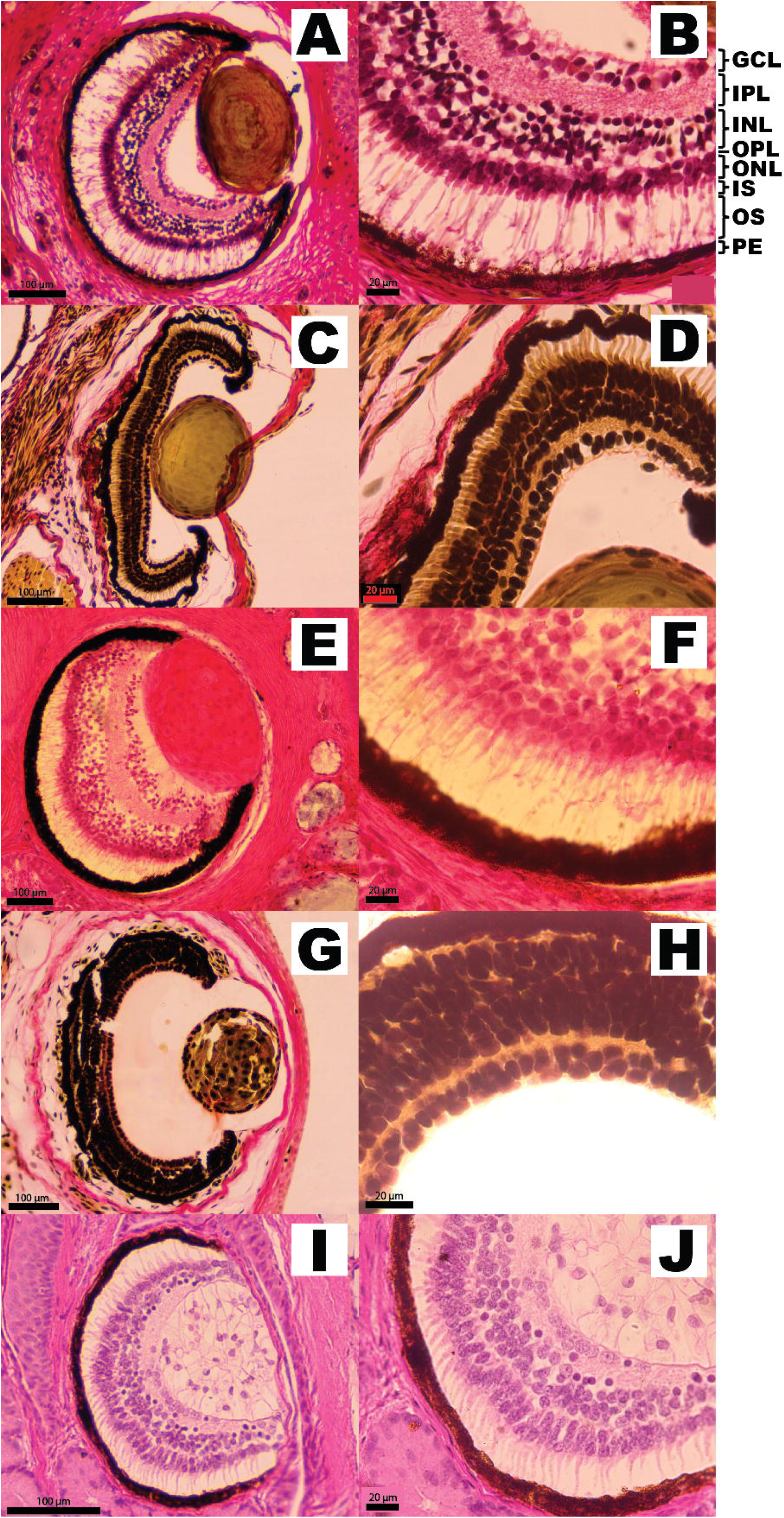
Photomicrographs demonstrating photoreceptor cell diversity. Black arrows point to photoreceptors that exhibit characteristics typical of cone photoreceptors (bulbous ellipsoids, tapered outer segments that are much thinner than their inner segments); blue arrow indicates a photoreceptor reminiscent of a double rod. These are morphologically distinct from the rods that predominate the caecilian retina. Beside each panel, we include diagrams and images of cone photoreceptors and one double rod diagram that resemble the structures we identified. (A) *Epicrionops sp.* (B) *Epicrionops sp.* (C) *Dermophis mexicanus* (D) *Ichthyophis kohtaoensis* (E) *Scolecomorphus kirkii* (F) *Dermophis mexicanus;* Dissociation of the photoreceptors from the pigment epithelium (PE) is a frequent artifact of preservation. Scale bars are 5 µm (A, B, C, D, E) and 20 µm (F). The retinal cell drawings are from (Walls, 1942). The blue diagrams were adapted from Donner and Yovanovich (2020). The grey images were adapted from Isayama et al. (2014). Abbreviations for the diagram associated with (B): o - outer segment; d - oil droplet; e - ellipsoid; m - myoid; l - external limiting membrane; n - nucleus; f - foot-piece.

**Table 1.** Summary of characters reported from sections shown in Figure 1.

| Species | Inner Rod Segment Length ($\mu\text{m}$) | Outer Rod Segment Length ($\mu\text{m}$) | Outer Plexiform Layer width ($\mu\text{m}$) | Inner Plexiform Layer width ($\mu\text{m}$) | Number of nuclei rows in ONL | Number of nuclei rows in INL | Number of nuclei rows in GCL | Lens Shape |
| --- | --- | --- | --- | --- | --- | --- | --- | --- |
| *Epicrionops* sp. | 10 | 35–40 | NA | 40 | 2–3 | 3–4 | 1–2 | ellipsoid |
| *Ichthyophis kohtaoensis* | 5 | 8–14 | 2–4 | 7–10 | 1–2 | 2–3 | 1–2 | ellipsoid |
| *Dermophis mexicanus* | 10 | 30–35 | NA | 46 | 2–3 | 3–4 | 2–3 | ellipsoid |
| *Typhlonectes compressicauda* | N/A<br>(see text) | N/A<br>(see text) | < 3 | 5–8 | 1–2 | 3–4 | 1–2 | spherical |
| *Scolecormorphus kirkii* | 8 | 15–20 | 2–5 | 12 | 2–3 | 2–3 | 1,<br>discontiguous | amorphous |

### 3.3 RETRIEVAL OF *LWS* FROM CAECILIANS

We retrieved *LWS* sequences from four species of Gymnophiona (*Rhinatrema bivittatum*, XM_029609805.1; *Microcaecilia unicolor,* XM_030192059.1; *Geotrypetes seraphini,* XM_033922369.1; *Ichthyophis kohtaoensis*, GCA_033557465.1). The sequences are publicly available in GenBank from Torres-Sanchez et al. (2019) and early results from the Vertebrate Genome Project ([https://vertebrategenomesproject.org/]; (Haussler et al., 2009)). Previous attempts to detect *LWS* were likely unsuccessful (Mohun et al., 2010) due to primer mismatch (see Supplementary Table 5 in section 3.3 of the Supplementary Material).

### 3.4 FURTHER VALIDATION OF A CAECILIAN *LWS* AND ASSESSMENT OF ITS POTENTIAL FUNCTIONALITY

#### 3.4.1 Transcriptomics and presence of LWS

In total, we obtained 131,623,768 reads from *C. orientalis* eye tissue (PRJNA1181370). The reads were assembled into a reference eye transcriptome containing 175,181 transcripts. For *RH1*, one transcript was assembled, while for *LWS*, two were assembled. Compared to LWS_v1, LWS_v2 is missing exon 5 which encodes transmembrane domains VI and VII as well as the C-terminal region, based on comparison with bovine rhodopsin. We found that *RH1* was expressed at much higher levels (17.53 TPM) compared to LWS_v1 (0.22 TPM) or LWS_v2 (0.02 TPM; 80.3X and 818.5X, respectively). We note that, given our expression data is based on a transcriptome of the whole eye, it does not necessarily reflect mRNA expression exclusive to the retina, though significant expression of *LWS* and *RH1* is not expected from extraretinal tissues of the eye. No sequences for *SWS1* or *SWS2* were assembled.

#### 3.4.2 PCR

We obtained 568 bp of *LWS* sequences from 11 species of caecilians. After aligning with *C. orientalis*, all sequences had open reading frames that lacked stop codons (Supplementary Data 6).

#### 3.4.3 λ_max_ of LWS and RH1

The estimated λ_max_ for the complete *LWS* gene sequences of five caecilian species (*G. seraphini*, *I. kohtaoensis*, *M. unicolor*, *R. bivittatum*, and *C. orientalis*) ranged from 508.57 to 550.58 nm. The shortest predicted absorbance value corresponds to *LWS*_v2 from *C. orientalis*. The predicted λ_max_values for *RH1* in caecilians, using the complete gene sequences, ranged from 492.07 to 513.54 nm (Figure 4; Supplementary Table 4).

**FIGURE 4.**
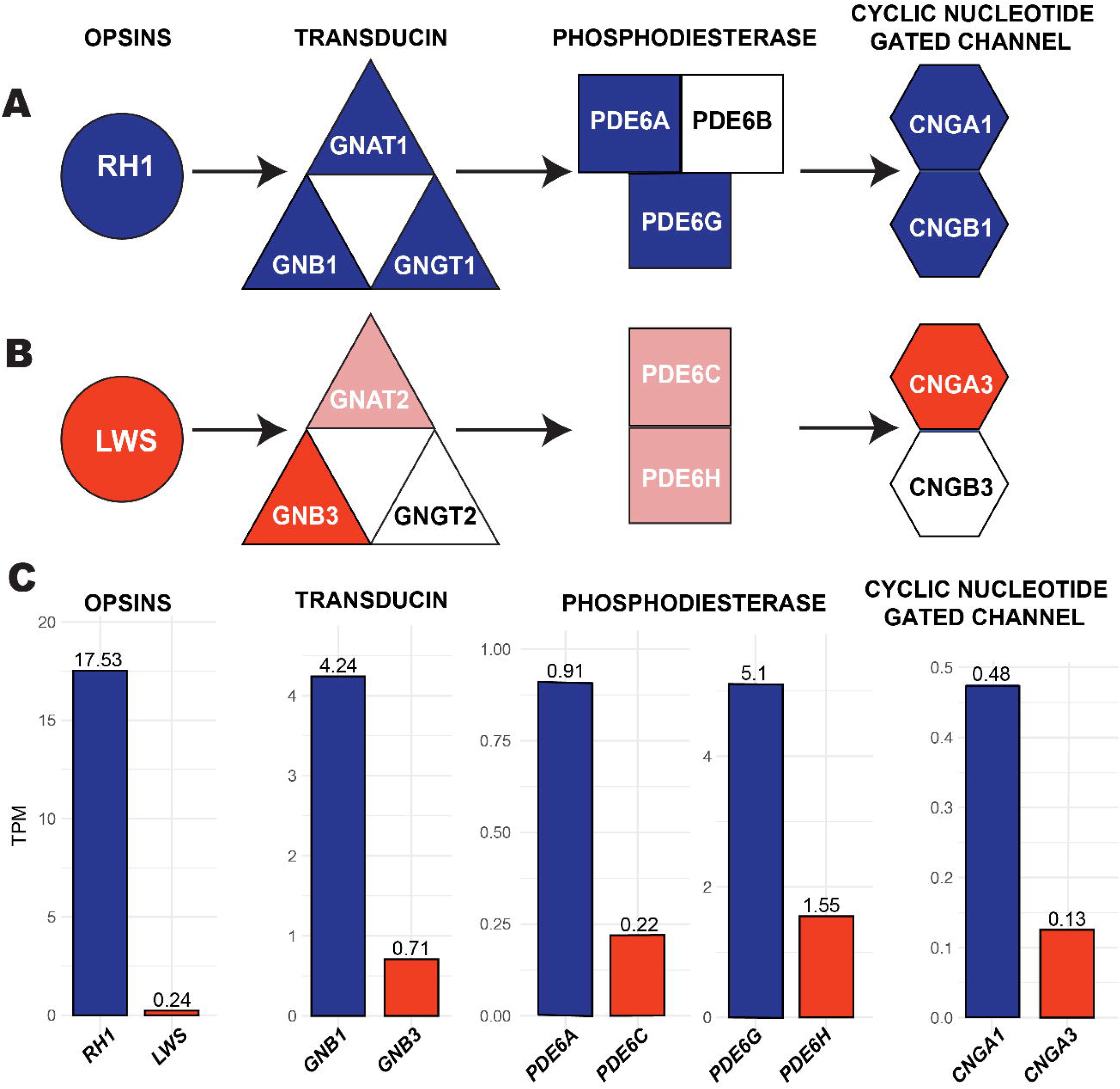
(A) Maximum wavelength absorbance values (λ_max_) for *LWS* and *RH1* estimated using a regression-based machine learning model (ML; solid markers; Frazer et al., 2024) or microspectrophotometry (MSP; hollow markers; Mohun et al., 2010) for five caecilian species. Because *LWS* was previously thought to be absent in caecilians, no MSP data is available for this opsin. The color scale at the top of the figure represents the visible light spectrum for the range of estimated absorbance values (470–570 nm). The image corresponds to *Microcaecilia albiceps*. (B) Linear regression between ML and MSP data for *LWS* (Pearson’s correlation test: r = 0.13 , P = 0.66) and *RH1* ( r = 0.38 , P = 0.09). The images show three caecilian species included in our study: *C. orientalis*, *M. albiceps* (© Santiago Ron; BIOWEB, https://bioweb.bio), and *G. seraphini* (© Brian Freiermuth, with permission).

We estimated absorbance values for *LWS* and *RH1* in 39 additional vertebrate species, with *LWS* values ranging from 531.39 nm in the lungfish *Protopterus annectens* to 565.3 nm in the chicken *Gallus gallus*, and *RH1* values ranging from 493.72 nm in the lizard *Pogona vitticeps* to 513.54 nm in the frog *Scaphiopus couchii* (Supplementary Table 4).

Of the 44 species we reviewed, microspectrophotometry (MSP) data were available for 14 *LWS* sequences and 21 *RH1* sequences, including for two caecilian species. Our results suggest a weak relationship between the λ_max_ values estimated using machine learning and experimental MSP data from the literature, particularly for *LWS* (Pearson’s correlation: □ = 0.13, □ = 0.66). Although not significant, the correlation is stronger for *RH1* (□ = 0.38, □ = 0.09; Figure 4B). Additionally, the machine-learning estimates showed greater discrepancies from MSP data for *LWS* (mean absolute error: 26.4 nm) compared to *RH1* (2.8 nm). For the two caecilian species with *RH1* MSP data, the difference between MSP and machine learning data was ∼3 nm (Figure 4A; Supplementary Table 4); MSP data for *LWS* was not available to compare.

#### 3.4.4 Assessing phototransduction cascade genes to infer LWS expression location

Our results indicate that most components of the rod phototransduction pathway are intact in caecilian genomes and are expressed in the eye transcriptome of *C. orientalis* (with only 1 of 8 genes absent in all species; Figure 5A, Table 2). In contrast, the cone pathway displays a more mosaic pattern, with 2 of 7 genes absent from all species and 3 others missing in one or more species (Figure 5B, Table 2).

**FIGURE 5.**
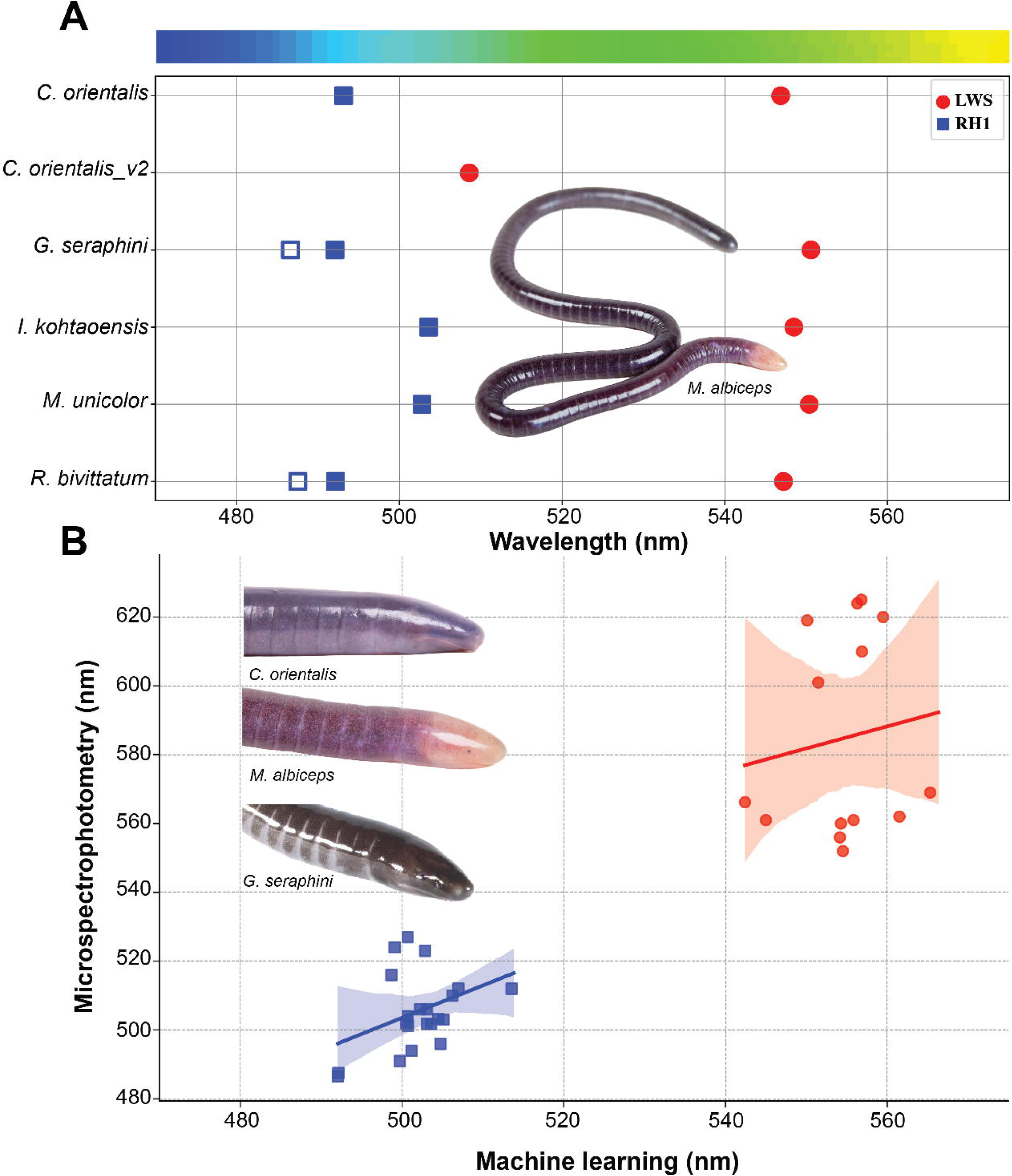
Rod and cone phototransduction cascade components and expression levels in caecilian eye tissue. (A) In blue, schematic of the rod phototransduction cascade, showing the key proteins involved: the rod opsin *RH1*, the heterotrimeric transducin complex (*GNAT1*, *GNB1*, *GNGT1*), phosphodiesterase components (*PDE6A*, *PDE6B*, *PDE6G*), and the cyclic nucleotide-gated channel subunits (*CNGA1*, *CNGB1*). (B) In red, schematic of the cone phototransduction cascade, highlighting the cone opsin *LWS* and its associated components: transducin complex (*GNAT2*, *GNB3*, *GNGT2*), phosphodiesterases (*PDE6C*, *PDE6H*), and cyclic nucleotide-gated channel subunits (*CNGA3*, *CNGB3*). (C) Bar plots showing transcript expression levels (transcripts per million, TPM) for the rod- and cone-associated phototransduction genes that are present in the eye transcriptome of *Caecilia orientalis.* Blue bars indicate rod-associated genes; red bars indicate cone-associated genes. White shapes in (A) and (B) indicate genes not detected in the genomes or transcriptome, while light blue and light red indicate genes present in at least one, but not all, of the assessed species (see Table 2).

**Table 2.** Presence and characteristics of rod and cone phototransduction cascade genes in four caecilian species. Symbols and cell colors indicate detection (+; black cell), absence (-; white cell), or presence with sequence irregularities (e.g., early stop codon, shorter or longer sequence, ambiguous sequence; grey cell).

| Protein complex | Gene Symbol | Cell type | *R. bivittatum* | *M. unicolor* | *G. seraphini* | *C. orientalis* transcript |
| --- | --- | --- | --- | --- | --- | --- |
| Transducin | *GNAT1* | Rod | Early stop codon | + | + | + |
|  | *GNB1* |  | + | + | + | + |
|  | *GNGT1* |  | + | + | + | + |
|  | *GNAT2* | Cone | + | - | - | - |
|  | *GNB3* |  | + | Variable sequence | + | + |
|  | *GNGT2* |  | - | - | - | - |
| Phosphodiesterase | *PDE6A* | Rod | + | + | + | Shorter sequence |
|  | *PDE6B* |  | - | - | - | Ambiguous sequence |
|  | *PDE6G* |  | + | + | + | + |
|  | *PDE6C* | Cone | + | - | + | Shorter sequence |
|  | *PDE6H* |  | + | - | - | Longer sequence |
| Cyclic Nucleotide Gated Channel | *CNGA1* | Rod | + | + | + | + |
|  | *CNGB1* |  | + | + | + | + |
|  | *CNGA3* | Cone | + | + | + | Shorter sequence |
|  | *CNGB3* |  | - | - | - | - |

Most genes are represented by two or more isoforms, though these may be assembly artifacts. We provide alignments including all detected transcriptomic isoforms in Supplementary Data 7.

Within the rod phototransduction pathway, *PDE6B*—a subunit of the phosphodiesterase complex—is the only gene absent from all available caecilian genomes and from the salamander *P. waltl*, suggesting a possible gene loss. A truncated *PDE6B*-like sequence (127 amino acids; typical length ∼890) was detected in the *C. orientalis* transcriptome (Figure 5A, Table 2), sharing 26.8% similarity with *C. orientalis PDE6A*. Due to its short length and absence in other caecilians, this annotation remains uncertain. For comparison, we included the putative *PDE6B* sequence in both *PDE6A* and *PDE6B* alignments (Supplementary Data 7). Although the remaining rod genes were present in caecilians, several exhibited sequence irregularities (Table 2), which are described in detail in Supplementary Material Results section 3.4.4.

In contrast, the cone phototransduction pathway shows more instances of gene loss and a mosaic pattern of gene presence and absence in caecilians (Figure 5B, Table 2). The genes *GNGT2* and *CNGB3*, which encode components of the cone transducin complex and the cyclic nucleotide-gated (CNG) channels respectively, are entirely absent from all available genomes and transcriptomes of Gymnophiona. Meanwhile *GNB3* and *CNGA3*, which are also part of the transducin complex and CNG channels, are the only cone phototransduction genes consistently present across all caecilian species, though both exhibit irregularities in *M. unicolor* and *C. orientalis*, respectively (Table 2). Additional details on cone phototransduction gene presence/absence and sequence irregularities are provided in Supplementary Material Results section 3.4.4.

In terms of expression levels of rod- versus cone-specific phototransduction genes in *C. orientalis*, the pattern observed mirrors that of the opsins, in that genes associated with the rod phototransduction pathway are expressed at higher levels than those associated with the cone phototransduction pathway (Figure 5C).

### 3.5 SELECTION ANALYSES

Results from the complete and incomplete *LWS* analyses were congruent (Supplementary Material File S1 and S2). Thus, we only present the complete dataset. Results from the incomplete dataset are provided in Supplementary Material Section 3.5.1. We initially tested whether purifying selection better described the data than neutral evolution, which would be consistent with long-term maintenance of *LWS*. In this test, the unconstrained M0 model was favored over the constrained M0 model (null hypothesis), in which omega was fixed at 1 (simulating neutral selection), for both *LWS* (likelihood ratio (LR) = 2687.642, P << 0.0001) and *RH1* (LR = 2244.112, P <<0.0001), rejecting the null hypothesis. Additionally, we found that the M3 model was a better fit than M0 for both genes (*RH1*: LR = 615.675, P < 0.0001; *LWS*: LR = 825.605, P < 0.0001; File S2), suggesting significant ω variation across sites. To test for positively selected sites, we compared the fit of the M8 model, which allows for sites with ω > 1, to that of M7 and M8a. For *RH1*, the M8 model was a better fit than M7 (LR = 40.181, P < 0.0001; File S2) and M8a (LR = 5.978, P = 0.015; File S2). Two sites were inferred to be under positive selection with a BEB posterior probability >90% (positions 213 and 217; bovine *RH1* numbering), while one site was close to but did not meet the significance threshold (position 107). Consistently, BUSTED found nearly significant evidence for gene-wide episodic diversifying selection of *RH1* in caecilians (P = 0.059) and identified two sites with evidence ratios (ER) ≥ 100 (positions 107 and 213, File S1), aligning with CODEML.

In contrast, for *LWS*, M8 was a better fit than M7 (LR = 44.241, P < 0.0001; File S2) but not M8a (LR = 0.0, P = 1.0; File S2), indicating only a weak signal for positive selection. Only one site exceeded the 90% BEB significance threshold (site 66; 49 in bovine *RH1*). Meanwhile, BUSTED did not detect gene-wide episodic diversifying selection in caecilian *LWS* sequences (P = 0.081) and found no sites under positive selection (File S1).

The FEL and FUBAR analyses can detect signals of positively selected sites across the entire phylogeny. However, FUBAR did not identify any sites for *LWS* or *RH1*. On the other hand, FEL identified one site under diversifying positive selection in *LWS* when the stem caecilian clade was used as foreground (site 109; 92 in bovine *RH1;* P = 0.04), but not across the entire phylogeny. In contrast, for *RH1*, FEL identified one site under positive selection across the entire phylogeny (119; P = 0.05) and a different site when the stem caecilian clade was analyzed as foreground (292; P = 0.04). Interestingly, FEL results showed that 272 out of 368 codon sites (when no foreground was identified) and 175 out of 368 codon sites (when caecilians were defined as the test group) in *LWS* were under purifying selection at p ≤ 0.05. This pattern of purifying selection is similar to that observed in the functional *RH1*, where 247 out of 354 codon sites (no foreground) and 165 out of 354 codon sites (caecilians foreground) were under purifying selection at the same significance threshold. These findings suggest that selection is acting to preserve the integrity of visual opsin gene sequences and maintain their function over evolutionary time.

Our analysis using RELAX revealed an overall relaxation of selective constraint in *LWS* but not *RH1* when crown Gymnophiona and its stem branch were designated as the foreground (*LWS:* K = 0.74, LR = 15.47, P < 0.0001; *RH1*: K = 1.37, LR = 1.67, P = 0.196; Supplementary Figure 1 and File S1). This pattern was further supported by CODEML. Specifically, both clade model C (CmC vs M2a_rel: LRT = 8.507, P = 0.004; File S2) and clade model D (CmD vs. M3: LRT = 11.055, P = 0.0009; File S2) were supported over other models, providing significant evidence for shifts in selection pressures, with some sites in *LWS* showing a transition from relatively stronger to weaker purifying selection within Gymnophiona + stem. For *RH1*, however, neither the CmC nor CmD models were favored over the alternative models (CmC vs M2a_rel: LRT = 0.508, P = 0.476; CmD vs M3: LRT = 3.053, P = 0.08), suggesting that selective pressure on *RH1* has not undergone significant evolutionary shifts in caecilians (File S2).

## 4. DISCUSSION

### 4.1 Evolution of caecilian eyes

Caecilians (Gymnophiona) are an ancient order of amphibians whose ancestor began transitioning to a fossorial lifestyle more than 250 million years ago (Kligman et al., 2023; Santos et al., 2020). Caecilians have undergone a degeneration of visual organs associated with an increasingly fossorial lifestyle (Himstedt, 1995; Kamei, Mauro, et al., 2012; Taylor, 1968), including losing two of three cone opsin genes (Lin et al., 2024; Mohun et al., 2010; Musser & Arendt, 2017). As evidenced by our ancestral state reconstructions of eye musculature and eye exposure (Figure 1, Supplementary Material Methods section 2.1), patterns in trait loss are not necessarily correlated with species ecology or phylogeny. For example, *Dermophis mexicanus* and *Gymnopis multiplicata* are closely related and ecologically similar members of Dermophiidae from Central America: they are viviparous, live as litter burrowers in shallow soil, and emerge at dawn and dusk, especially in light rains, to forage (MHW pers. obs.). *Dermophis mexicanus* have relatively well-developed eyes in open sockets, covered by skin, with all six extrinsic muscles, a cell-rich, organized retina, and a crystalline lens that retains some cells peripherally. In contrast, *Gymnopis multiplicata* has eyes in sockets covered by bone as well as skin, no extrinsic muscles, and an amorphous cellular lens that is usually flattened against the cornea and retina (Wake, 1985). In another example, the covering of the eye socket by bone is a homoplastic character that occurs in members of several families: Caeciliidae (*Oscaecilia* and *Caecilia*), Dermophiidae (*Gymnopis*, *Dermophis*), Herpelidae (*Boulengerula*), and Scolecomorphidae (*Scolecomorphus*, *Crotaphatrema*). *Scolecomorphus* has an exceptional morphology: the eyes in at least three species are not in the bone-covered sockets but embedded in the tentacle (De Jager, 1940; Taylor, 1968; Wake, 1985) see also Supplementary Material section 4.3 (De Jager, 1940; Taylor, 1968; Wake, 1985). Given what appears to be a general association between a bone-covered socket and reduced eye musculature (Figure 1), it is tempting to consider the covered eye as a factor driving further degeneration of the visual apparatus in these taxa (Wake, 1985), yet the morphological patterns are not exactly similar, with different elements being more reduced or lost among the species in the set, including among *Scolecomorphus* species. These examples illustrate the mosaic loss of visual features across the history of caecilians, likely because of the relaxed selection that appears to be a feature of the clade.

### 4.2 Expression of LWS, phototransduction cascade genes, and morphology of caecilian retinae

For such a unique and ancient vertebrate order, caecilians are woefully understudied, and their genomes are no exception (Torres-Sánchez et al., 2019). Prior to work by Lin et al (2024), caecilian eyes were thought to have rod-based retinae that only contain rhodopsin (Mohun et al., 2010; Wake, 1985). Here we report full-length transcriptomic *LWS* and *RH1* sequences from one additional caecilian family (Caecilidae), as well as partial *LWS* sequences from four additional families (Caecilidae, Herpelidae, Typhlonectidae, and Scolecomorphidae), expanding coverage to eight of the ten families in the Order Gymnophiona. These findings indicate that *LWS* has likely been maintained across all extant lineages, despite varying degrees of fossoriality and eye reduction.

Given that all these *LWS* sequences have a translatable open reading frame and a stop codon position homologous to other amphibian *LWS* sequences (Supplementary Data 3), and that the gene is expressed in the eye of at least one caecilian species (Figure 5C), we hypothesize that *LWS* is functionally expressed. The maintenance of *LWS* is further supported by our selection analyses, which indicates that although there is a signal of relaxed evolution, a model of purifying selection provides a significantly better fit than a fully neutral model—unsurprising given the gene’s retention over 200 million years of fossorial evolution. However, results from two complementary approaches (CmC and CmD, and RELAX) indicate that *LWS* in caecilians is experiencing a relaxation in the strength of purifying selection relative to other amphibians. This pattern is not observed in the other retained opsin, *RH1*, and may suggest that the physiological role of *LWS* is under less stringent selective constraint. We explore this possibility further in the next section.

Our *LWS* expression data comes from *Caecilia orientalis*, a member of Caeciliidae known for derived fossorial adaptations and relatively reduced eyes (*Taylor 1968*, n.d.). In this species, *LWS* expression is significantly lower than *RH1*. There is little eye transcriptome data available for amphibians but these results align with data from the nocturnal frog *Rana sphenocephala*, where adult individuals express *RH1* at about 50 times the level of *LWS* (Schott et al., 2022). Similarly, in some dim-light adapted (e.g., deep-water, nocturnal) fish species, *LWS* expression relative to *RH1* can be very low (Fogg et al., 2022; Luehrmann et al., 2019).

However, some diurnal species also show relatively low *LWS* expression, such as the spotted unicorn fish (*Naso brevirostris*) (Tettamanti et al., 2019). The relatively low expression of *LWS* may be a result of a rod-biased (or rod-only) retina, a pattern observed across tetrapods, but especially pronounced in nocturnal lineages or those that were adapted to nocturnality at some point in their evolutionary history (de Busserolles et al., 2021; Kim et al., 2016; even at relatively young evolutionary time scales, e.g., in colubrid snakes; Walls, 1942). The common ancestor of Amphibia was likely nocturnal (Anderson & Wiens, 2017), and salamanders and anurans possess a higher proportion of rods to cones in their retinae (Donner & Yovanovich, 2020; Fain, 1976; Nilsson, 1964; Walls, 1942).

While the presence of intact and expressed *LWS* and *RH1* genes suggests the possibility of a duplex retina, opsin expression alone does not confirm the maintenance of two functionally distinct photoreceptor cell types. To further test this hypothesis, we pursued two complementary lines of evidence: we examined the presence and expression of key rod and cone phototransduction cascade genes and evaluated the morphology of caecilian retinas for structures resembling cones. Our results revealed that most components of the rod phototransduction pathway are present in caecilian genomes and expressed in the eye of *C. orientalis* (Figure 5A), supporting the retention of rod function. In contrast, we observed a mosaic of gene loss in the cone phototransduction pathway, consistent with relaxed selection on color vision in caecilians (Figure 5B). The only rod-specific gene we were unable to detect in any caecilian genome was *PDE6B*. This gene encodes one of the catalytic subunits of the PDE6 holoenzyme, which is essential for signal transmission in rods as it hydrolyzes cGMP, leading to the closure of cyclic nucleotide-gated channels and subsequent membrane hyperpolarization (Lagman et al., 2016).

Notably, *PDE6B*, *PDE6A*, and *PDE6G* have also been reported as lost in nocturnal geckos (Pinto et al., 2019; Schott et al., 2019). Instead, these species are thought to utilize the cone protein PDE6C to transduce rod signaling, supporting the hypothesis that ancestral cones were transmuted into rods for scotopic vision (Kojima et al., 2021; Zhang et al., 2006). Similarly, *PDE6A* is lost in birds and reptiles (Lagman et al., 2016), where this loss may be compensated by the formation of a *PDE6B/B* homodimer rather than the *PDE6A*/*B* heterodimer typically used by other vertebrates (Huang et al., 2004). As such, the absence of *PDE6B* in caecilian genomes does not necessarily imply that the rod pathway is non-functional; instead, it raises the possibility that an alternative catalytic subunit—potentially even *PDE6C*—may compensate for cGMP hydrolysis in caecilian rods. It is important to note, however, that *M. unicolor* appears to lack both *PDE6B* and *PDE6C*, though this observation could reflect incomplete sequencing or assembly artifacts. Alternatively, if these absences are real, they raise further questions about how cGMP hydrolysis is achieved in this species.

In contrast, cone phototransduction genes exhibit a mosaic pattern of presence and absence across caecilian species. The only cone phototransduction gene consistently present in all species analyzed is the Guanine nucleotide-binding protein β3 (*GNB3*) (5B, Table 3).

However, in addition to its expression in cone photoreceptors, *GNB3* has also been reported in subsets of bipolar cells and rod photoreceptors across vertebrates, so it is possible that the transcripts we detect originate from these other retinal cell types (Dhingra et al., 2012; Ritchey et al., 2010). Many critical components are missing in all four surveyed species or detected only in one or, at most, two caecilian species (Figure 5B, Table 3). The potential impact of these gene absences in caecilians can be inferred from experimental studies in mice. For example, the absence of *CNGA3* and *CNGB3* leads to cone degeneration, affecting both cell viability and function (Michalakis et al., 2010). Moreover, although the effects of *GNGT2* loss have not been directly studied, findings from *GNAT2* suggest that any disruption to the transducin complex would likely impair cone phototransduction without necessarily compromising cone cell survival (Chang et al., 2006; Ronning et al., 2018).

A similar patchy pattern of cone phototransduction gene losses has been reported in other species that are either adapted to low-light conditions or with a history of subterranean lifestyles during their evolution. For example, one species of armadillo (*Dasypus novemcinctus*) lacks *CNGB3* (Emerling & Springer, 2015) and one sloth species (*Choloepus hoffmanni*) and a group of subterranean snakes lack *GNGT2* (Emerling & Springer, 2015; Gower et al., 2021). These losses, along with the inactivation of other cone-specific genes, have been interpreted as evidence for rod monochromacy in these groups. Given the extent of cone phototransduction gene loss in Gymnophiona—with all species lacking at least two of the seven genes examined, and in some cases, such as *M. unicolor*, lacking five genes—it is tempting to speculate that caecilians may lack functional cones altogether, resulting in a rod-only retina. Taken together, our findings suggest that while caecilians have retained *LWS*, their cone phototransduction machinery may be at least partially degraded. This raises the possibility that *LWS* is expressed in rod photoreceptors, and that some cone phototransduction components may have been co-opted in rod cells to support *RH1* function and/or some rod phototransduction components may enable *LWS* functionality.

Furthermore, existing literature and our morphological review of caecilian retinae do not provide conclusive support for the presence of cone cells in any caecilian species. Cone cells could be overlooked if they are inconspicuous and highly outnumbered by large rod cells, which is consistent with our 80:1 *RH1* to *LWS* mRNA transcript ratio. Thus, observations of photoreceptors with cone-like characteristics could be dismissed as anomalous variation, damaged cells, or rod outer segments in a state of regeneration. We carefully inspected over 280 images in order to encounter a handful of putative cone-like cells. There is considerable morphological variation among these cone-like photoreceptors, and, given the extreme variation in cone morphology observed among vertebrates (even at relatively young evolutionary time scales, e.g., in colubrid snakes; Walls, 1942); it is impossible to know what caecilian cones might look like and how similar or different they might be among families, some of which likely diverged nearly 200 MYA (Kligman et al., 2023). We acknowledge that given the distortions and artifacts observed in retinal sections, the cone-like photoreceptors we highlighted may be rod cells with anomalous or misleading morphologies as described above.

The expression of cone opsins in rod photoreceptors, or the co-expression of multiple visual opsins within a single photoreceptor cell, is not uncommon. For example, the cone opsin *SWS2* is expressed in ‘green rods’ of frogs and salamanders (W. I. L. Davies et al., 2012), which, in conjunction with ‘red rods’ expressing *RH1,* is thought to aid in nocturnal color vision (Yovanovich et al., 2017). Additionally, Gower et al. (2021) found that *LWS* was co-expressed with *RH1* in rod cells of *Anilios bicolor,* a fossorial scolecophidian snake with highly reduced eyes. Thus, a similar phenomenon could be occurring in caecilians. Immunolabeling studies employing antibodies against *LWS* or other proteins specific to cones and the cone phototransduction pathway will be necessary to confirm their presence in the caecilian retina.

As far as we are aware, our study presents the first set of eye transcriptome data for any caecilian, so we are not certain if the two *LWS* transcripts that we identified are artifacts of the assembly process or represent alternatively spliced transcripts. The genome assemblies of *R. bivittatum*, *M. unicolor*, and *G. seraphini* only contain one annotated transcript, but as far as we are aware, the assembly process did not incorporate eye transcriptome data (Ovchinnikov et al., 2023). Based on our alignments with bovine rhodopsin, exon 5 encodes portions of transmembrane domains VI and VII and the C-terminal region of the opsin protein—regions that play essential roles in chromophore binding (Palczewski et al., 2000) and spectral tunning .

Therefore, the absence of exon 5 in one of the *LWS* transcripts likely renders that isoform non-functional, at least in its expected purpose.

### 4.3 Evidence for and implications of a putatively functional LWS in caecilians

When analyzing the amino acid composition of known spectral tuning sites in available *LWS* and *RH1* sequences for caecilians, we found that they are consistent with data reported for other amphibians (Schott et al., 2024; Wan et al., 2023), containing at least one of the known amino acid variants at each site (see Supplementary Table 6 in section 4.2). For example, as highlighted by Lin et al. (2024) the specific combination of S164, H181, Y261, T269, and A292 found in the *LWS* sequence of *R. bivitatum*, *G. seraphini*, *M. unicolor,* and *C. orientalis* has been shown to generate a long-wavelength-sensitive photopigment with λ_max_at ∼560 nm in other vertebrates (Asenjo et al., 1994; Hauzman et al., 2017; Yokoyama, 2002b; Yokoyama & Radlwimmer, 2001b). While this prediction is relatively close to our machine learning estimates (546.87– 550.58 nm; excluding *C. orientalis LWS*_v2), our estimates fall on the shorter end of the whole range of absorption observed in the vertebrate *LWS* (∼549–626 nm; Schott et al., 2024), similar to the absorbance properties observed in nocturnal terrestrial species (Margetts et al., 2024; Veilleux & Cummings, 2012). If accurate, these short absorbance values could enhance low-light vision by improving the signal-to-noise ratio through reduced thermal noise susceptibility (Luo et al., 2011).

It is important to note, however, that our machine learning (ML) model showed only a weak correlation with MSP data for *LWS* ( = 0.13, = 0.66), possibly because of an underrepresentation of opsins in the ∼600-nm range within the ML training dataset (Frazer et al., 2024). Moreover, the model was trained using absorbance values based on 11-cis retinal (A1) as the chromophore (Frazer et al., 2024), which may result in estimates that differ from MSP values obtained using alternative chromophores such as A2. Additionally, variation in MSP data cannot be ruled out; for instance, red cones from *Ambystoma tigrinum* show a 30–40 nm range in MSP data (Makino & Dodd, 1996). This variation likely stems from the challenges of estimating absorbance in single cone photoreceptors, especially for *LWS* in amphibians, as it is more susceptible to noise and generally has smaller sample sizes compared to rod pigments like *RH1* or *SWS2* (see Supplementary material in Schott et al., 2024).

*LWS* is present in a significant proportion of fossorial and low-light adapted taxa, such as blind mole rats, cryptozooic Australian scincid lizards, and some deep-sea fish species (W. I. L. Davies et al., 2012; Ford et al., 2024; Musilova et al., 2021). Most reported instances of *LWS* gene loss or pseudogenization occur in fishes adapted to deep sea environments (Lin et al., 2017; Musilova et al., 2021), where light in the long wavelength end of the visual spectrum is minimal or absent. To our knowledge, pseudogenization of *LWS* in tetrapods has only been observed in the subterranean naked mole-rat (*Heterocephalus glaber*), the Cape golden mole (*Chrysochloris asiatica*), armadillos, and deep diving whales (Emerling & Springer, 2014; Meredith et al., 2013). Dysfunction or complete loss of *LWS* is possible in the obligate subterranean Texas blind salamander (*E. rathbuni*), given that Tovar et al. (2021) were unable to label any of the visual opsins in the retina. However, failure to immunochemically label LWS in the retina is not necessarily confirmation of gene loss or dysfunction. For example, in one population of the blind cave salamander (*Proteus anguinus anguinus*), Kos et al. (2001) found the LWS protein in the pineal gland but not the eye, while both tissues from other populations were successfully immunolabeled for LWS (Kos et al., 2001). These data show that the *LWS* gene tends to persist in ecological and evolutionary contexts under which other cone opsins are readily lost, strongly suggesting that *LWS* retains some function even in species with highly reduced visual systems.

Additional evidence for the potential functionality of *LWS* in caecilians arises from our selection analyses. With the shift to a fossorial lifestyle, caecilians underwent a simplification of their visual systems, likely leading to relaxed selection on the molecular mechanisms underlying vision, and eventually to the loss of some cone opsins and phototransduction genes. While the loss of *SWS1* and *SWS2* (Lin et al., 2024; Mohun et al., 2010) might be the result of relaxed selection, the same selective environment did not result in the loss of *LWS*, although the signal of relaxation remains detectable in caecilian *LWS* sequences (see Results section 3.3). Importantly, this relaxation in selective pressure did not result in an overabundance of deleterious mutations, as the *LWS* sequences retrieved from caecilian genomes and transcriptome have intact open reading frames. This contrasts with the other cone phototransduction genes that we examined, where 5 of 7 appear lost in at least one species, several are truncated in the *C. orientalis* transcriptome, and those that are variably present are always present in the earliest branching lineage, *R. bivittatum* (Table 3). Given this long-term retention of *LWS* (over 240 million years; Kligman et al., 2023), we hypothesize that some degree of purifying selection has preserved its functionality, despite the relaxed constraints associated with reduced visual demands. One possible function for *LWS* could be tied to behavioral ecology. Light between 520 and 610 nm is most prominent in the forest understory while in more open woodland environments wavelengths below 500 nm are more prevalent (Endler, 1993). Thus, slight variations in blue to red wavelengths may provide caecilians information about canopy cover, habitat type, and exposure to predators when they are at or near the surface in forested habitats. Furthermore, there are significant shifts in light intensity within forests at long and short wavelengths in the transition from daylight, to sunset, to twilight (Endler, 1993). Dichromatic vision is possible with only rods (*RH1*) and red cones (*LWS*) under mesopic conditions (W. I. L. Davies et al., 2012; Hunt et al., 2009) and could afford caecilians the spectral resolution necessary to discriminate between these structural and temporal variables of their environment. Alternatively, If *LWS* is expressed in rods in caecilians, similar information may be available to them at dawn/dusk/twilight and while moving in the leaf litter or just under the surface. As caecilians appear to be primarily nocturnal (Prakash et al., 2024) and most observations of surface activity occur at night, twilight or after heavy rains, when flooded underground habitats force animals to the surface to breathe, distinguishing light from different ends of the spectrum may be important for informing activity patterns, entraining circadian rhythm, and survival. If expressed in cones, *LWS* may also provide some visual information when caecilians find themselves at the surface under bright daylight conditions that would render rods useless due to saturation. Light intensity and spectra may be the only visual information available to caecilians. Therefore, retaining some spectral discrimination may allow them to maximize the visual information available given the constraints imposed by adaptation for head-first burrowing.

## 5. CONCLUSION

Although previous work suggested that the loss of *LWS* might be a synapomorphy of caecilians, our results show that these amphibians retain the *LWS* gene along with most rod-specific phototransduction genes and a subset of cone-pathway components. While it is possible that *LWS* is expressed in rod photoreceptors, its precise function remains uncertain. Selection analyses indicate that *LWS* is broadly under purifying selection, but with signs of relaxed constraint.

Although purifying selection appears strong enough to preserve gene integrity, this relaxation may reflect reduced reliance on canonical cone-mediated vision. Additionally, we detected *LWS* expression in the eye of one caecilian species but cannot confirm whether it is translated into a functional protein. The maintenance of the open reading frame, however, suggests some function of the mRNA, and/or of the protein, if it exists. We propose that *LWS* may function to transmit light signals to other genes that regulate activity patterns (Boyette et al., 2024) or that it is used to complement wavelength or luminance information obtained with *RH1.* Although traits are almost never completely lost (Sadier et al., 2022), it is interesting that *LWS*, which is a terminal gene in the visual cascade, has been maintained for such a long time under fossorial conditions. The maintenance of *LWS* in caecilians and other dim-light-adapted animals begs the question of what this opsin does and why it is still useful – i.e., why *LWS* and not the other opsins? These data pose new questions regarding the selective pressures associated with dim-light or fossorial lifestyles, the biology of caecilians, including the presence of bright coloration and above-ground behavior in some species, as well as present an interesting example of the mosaic nature of trait loss.

## Supporting information

FileS1_HYPHY_summary

FileS2_PAML_summary

## Author contributions

RDT, MJNM, SSA, JCS, SRR conceptualized the study; RDT, MJNM, SRR, SSA, JCS, MHW, JS collected data; MJNM, RDT, SSA, SSR performed analyses; MJNM, SSA, RDT, MHW, JS, JCS, SSR performed the investigation; RDT, SRR, JCS acquired funding; SRR, MJNM, MHW contributed samples and helped with permits; MJNM, SSA, JS, MHW, JCS, RDT, SSR prepared the original draft; RDT coordinated and supervised the project; all authors reviewed and revised the manuscript.

## Acknowledgments

We thank Rayna Bell and Lauren Sheinberg at the California Academy of Sciences, Carol Spencer at the Museum of Vertebrate Zoology, and Fernando Ayala and Diego Paucar at QCAZ for facilitating access to caecilian tissues for sequencing. We thank José Simbaña and Yanayacu Biological Station for permission and facilitation of specimen collection, Alexander Stubbs and Diego Paucar for their assistance in caecilian dissections and specimen preparation, Todd Oakley and Seth Frazer for their support in running the machine learning models to estimate λ_max_values for our sampled genes, Alex Somlyay for assistance with microscopy, Mike Boots for lending us use of his microscope and camera, Valeria Ramírez-Castañeda and Lydia Smith for conducting PCRs on MVZ tissues, Claudia Terán for laboratory work at QCAZ in Quito. Yocelyn Gutiérrez-Guerrero provided advice and guidance in the transcriptome analysis. For specimen collection in Ecuador, we thank William Booker, Verónica Crespo, Juan Guayasamin, Giovanna Romero, Yerka Sagredo, Scott Tiegs, and David Velalcazar. We thank Ryan Schott for his guidance on selection analyses, Simon Scarpetta, the two anonymous reviewers and the editor for their suggestions that improved the manuscript.

## Data accessibility statement

Sequences have been deposited in GenBank under accession numbers PQ541071–PQ541084 and in the Sequence Read Archive under project number PRJNA1181370. Voucher specimens are housed in the Museo de Zoología (QCAZ) at the Pontificia Universidad Católica del Ecuador, the Museum of Vertebrate Zoology (MVZ) at UC Berkeley, and the Herpetology Collection of the California Academy of Sciences. All other data are provided as supplementary material. Supplementary datasets have been updated and archived in Dryad (DOI: 10.5061/dryad.h18931zxf)

