## Supplementary material for "Caecilians maintain a functional long-wavelength-sensitive cone opsin gene despite signatures of relaxed selection and more than 200 million years of fossoriality"

### **Section 1- DATA**

**Supplementary Table 1 -** Character states and references for ancestral character reconstruction. Eye exposure: (0) eye exposed, (1) eye under skin, (2) eye under bone. Eye musculature: (0) all muscles lost, (1) two muscles, (2) three muscles, (3) four muscles, (4) six muscles, s. o. attenuate, (5) six muscles

| **Species** | **Eye exposure** | **Eye Musculature** | | **Source** | |
| --- | --- | --- | --- | --- | --- |
| *Boulengerula cf. boulengeri* | 2 | | 0 | | Wake 1993 |
| *Boulengerula taitanus* | 2 | | 0 | | Wake 1993 |
| *Caecilia gracilis* | 2 | | 1 | | Maciel and Hoogmoed 2011; Norris and Hughes 1913 |
| *Caecilia tentaculata* | 1 | | - | | Taylor 1968 |
| *Caecilia volcani* | 1 | | - | | Taylor 1968 |
| *Chthonerpeton indistinctum* | 1 | | 3 | | Wake 1993 |
| *Dermophis mexicanus* | 1 | | 5 | | Wake 1993 |
| *Epicrionops cf. marmoratus* | 1 | | 5 | | Taylor 1968 |
| *Geotrypetes seraphini* | 1 | | 3 | | Wake 1993 |
| *Gymnopis multiplicata* | 2 | | 0 | | Wake 1993 |
| *Hypogeophis rostratus* | 1 | | 5 | | Wake 1993 |
| *Ichthyophis* *kohtaoensis* | - | | 5 | | Wake 1993 |
| *Ichthyophis glutinosus* | 1 | | 5 | | Wake 1993 |
| *Idiocranium cf. russeli* | 1 | | 2 | | Wake 1993 |
| *Oscaecilia ochrocephala* | 2 | | 1 | | Wake 1993 |
| *Schistometopum thomense* | 2 | | 2 | | Wake 1993 |
| *Scolecomorphus vittatus* | 1 | | 1 | | Wake 1993; Wilkinson 1997 |
| *Typhlonectes natans* | 1 | | 4 | | Taylor 1968 |
| *Uraeotyphlus gansi* | 1 | | 5 | | Wake 1993 |

**Supplementary Table 2 - Examined specimens for morphological characterization of caecilian eyes.** Species IDs, taxonomic classification, S-book and museum numbers, along with figure and slide numbers, are provided for all specimens examined in the characterization of caecilian retinae and eye morphology. The examined slides correspond to archival specimens accessioned in the MVZ, originally prepared by MHW in the 1980s.

| **Family** | **Genus** | **Species** | **Ecology** | **S-Book** | **Field #** | **Museum #** | **Figure #: Slide # in Data 7** |
| --- | --- | --- | --- | --- | --- | --- | --- |
| Herpelidae | *Boulengerula* | *boulengeri* | Terrestrial burrower | 239 | 9004 | NA | NA |
| Caeciliidae | *Caecilia* | *occidentalis* | Terrestrial burrower | 185 | 318 | NA | NA |
| Typhlonectidae | *Chthonerpeton* | *indistinctum* | semi-aquatic burrower | 275F | NA | RDS 108 |  |
| Dermophiidae | *Dermophis* | *mexicanus* | Terrestrial burrower | 119 | 858 | NA | NA |
| Dermophiidae | *Dermophis* | *mexicanus* | Terrestrial burrower | 120 | 851 | NA | NA |
| Dermophiidae | *Dermophis* | *mexicanus* | Terrestrial burrower | 122 | 827 | NA | NA |
| Dermophiidae | *Dermophis* | *mexicanus* | Terrestrial burrower | 162 | 738 | NA | Fig. 3: 8  Fig. 4: 8, 9 |
| Dermophiidae | *Dermophis* | *mexicanus* | Terrestrial burrower | 164 | 754 | NA | NA |
| Dermophiidae | *Dermophis* | *mexicanus* | Terrestrial burrower | 172 | 1069 | NA | NA |
| Dermophiidae | *Dermophis* | *mexicanus* | Terrestrial burrower | 208 | 849 | NA | NA |
| Dermophiidae | *Dermophis* | *mexicanus* | Terrestrial burrower | 255B | 785 | MVZ 178976 | NA |
| Rhinatrematidae | *Epicrionops* | sp. | Terrestrial burrower | 367b |  | 27254 | Fig. 3: 98  Fig. 4: 97, 98 |
| Dermophiidae | *Geotrypetes* | sp. | Terrestrial burrower | 99 | 560 | NA | NA |
| Dermophiidae | *Geotrypetes* | *seraphini* | Terrestrial burrower | 168 | 1350 | NA | NA |
| Dermophiidae | *Gymnopis* | *multiplicata* | Terrestrial burrower | 166 | 2783 | NA | NA |
| Dermophiidae | *Gymnopis* | *multiplicata* | Terrestrial burrower | 167 | 131 | NA | NA |
| Grandisoniidae | *Hypogeophis* | *rostratus* | Terrestrial burrower | 183 | 428 | NA | NA |
| Grandisoniidae | *Hypogeophis* | *rostratus* | Terrestrial burrower | 450 | 1989.2079 | NA | NA |
| Ichthyophiidae | *Ichthyophis* | *kohtaoensis* | Terrestrial burrower | 448 | NA | NA | Fig. 3: 11  Fig. 4: 11 |
| Scolecomorphidae | *Scolecomorphus* | *kirkii* | Terrestrial burrower | 230L | NA | MCZ 27115 | Fig. 4: 30 |
| Scolecomorphidae | *Scolecomorphus* | *kirkii* | Terrestrial burrower | 230R | NA | MCZ 27115 | Fig. 3: 19 |
| Scolecomorphidae | *Scolecomorphus* | *uluguruensis* | Terrestrial burrower | 236 | NA | 12226 | NA |
| Typhlonectidae | *Typhlonectes* | *compressicauda* | Aquatic burrower | 101 | 836 | NA | NA |
| Typhlonectidae | *Typhlonectes* | *compressicauda* | Aquatic burrower | 171 | 134-10 | NA | NA |
| Typhlonectidae | *Typhlonectes* | *compressicauda* | Aquatic burrower | 241 | NA | NA | Fig. 3: 20 |
| Ichthyophiidae | *Uraeotyphlus* | *narayani* | Terrestrial burrower | 184 | iii | NA | NA |

#### **Supplementary Table 3 - Species sampling and associated information.** Species IDs, taxonomical classification, loci information, and NCBI accession numbers for all samples used in our selection analysis, maximum wavelength absorbance values (λ_max_) estimations, and synteny analysis. Museum numbers are included only for a limited subset of samples with available information.

| **Species** | **Order** | **Locus** | **Accession number** | **Museum number** |
| --- | --- | --- | --- | --- |
| *Ambystoma mexicanum* | Caudata | LWS | TSA: GFZP01130157 | NA |
|  |  | RHO | PGSH00000000 | NA |
| *Ambystoma tigrinum* | Caudata | LWS | AF038947 | NA |
|  |  | RHO | U36574 | NA |
| *Bombina bombina* | Anura | LWS | XM053696216 | NA |
|  |  | RHO | XM053688960 | NA |
| *Bombina pachypus* | Anura | LWS | TSA: GJVZ01067396 | NA |
| *Boulengerula boulengeri* | Gymnophiona | LWS | PQ541084 | CAS:Herp:168574 |
| *Bufo bufo* | Anura | LWS | TSA: GJPU01048513 | NA |
|  |  | RHO | U59921 | NA |
| *Caecilia orientalis* | Gymnophiona | LWS | PQ541083 | QCAZ:A:77338 |
|  |  | Transcriptome | PRJNA1181370 |  |
| *Chioglossa lusitanica* | Caudata | LWS | SRX7755094 | NA |
| *Cynops pyrrhogaster* | Caudata | LWS | AB043891 | NA |
|  |  | RHO | AB043890 | NA |
| *Dermophis mexicanus* | Gymnophiona | LWS | PQ541077 | MVZ:Herp:264229 |
| *Dermophis mexicanus* | Gymnophiona | LWS | PQ541082 | MVZ:Herp:263994 |
| *Epicrionops bicolor* | Gymnophiona | LWS | PQ541071 | QCAZ:A:11861 |
| *Epicrionops petersi* | Gymnophiona | LWS | PQ541072 | QCAZ:A:42405 |
| *Geotrypetes seraphini* | Gymnophiona | LWS | XM_033922369.1 | NA |
|  |  | RH1 | XP_033782390.1 |  |
| *Geotrypetes seraphini* | Gymnophiona | LWS | PQ541074 | MVZ:Herp:252475 |
| *Gymnopis multiplicata* | Gymnophiona | LWS | PQ541078 | MVZ:Herp:171331 |
| *Gymnopis multiplicata* | Gymnophiona | LWS | PQ541079 | MVZ:Herp:179536 |
| *Hyla sarda* | Anura | LWS | TSA: HBZV010637123 | NA |
|  |  | RHO | XM056528326 | NA |
| *Hynobius chinensis* | Caudata | LWS | TSA: GAQK01080456 | NA |
|  |  | RHO | TSA: GAQK01000060 | NA |
| *Hynobius retardatus* | Caudata | LWS | TSA: LE195300 | NA |
|  |  | RHO | TSA: LE184227 | NA |
| *Ichthyophis bannanicus* | Gymnophiona | Genome | GCA_033557465 | NA |
| *Ichthyophis bannanicus* | Gymnophiona | LWS | PQ541076 | MVZ:Herp:226266 |
| *Limnodynastes peronii* | Anura | LWS | SRR8712701 | NA |
|  |  | RHO | SRX5507031 | NA |
| *Microcaecilia albiceps* | Gymnophiona | LWS | PQ541075 | QCAZ:A:61704 |
| *Microcaecilia unicolor* | Gymnophiona | LWS | XM_030192059.1 | NA |
|  |  | RH1 | XP_030062458.1 | NA |
| *Microhyla fissipes* | Anura | LWS | TSA: GECV01009402 | NA |
|  |  | RHO | TSA: GECV01012731 | NA |
| *Oophaga pumilio* | Anura | LWS | TSA: GIKS01640506 | NA |
|  |  | RHO | SRR8275033 | NA |
| *Pelobates cultripes* | Anura | LWS | TSA: GHBH01058573 | NA |
|  |  | RHO | TSA: GHBH01063309 | NA |
| *Pyxicephalus adspersus* | Anura | LWS | TSA: ICLD02085713 | NA |
|  |  | RHO | TSA: PZQJ01000008 | NA |
| *Quasipaa spinosa* | Anura | LWS | SRX17159989 | NA |
| *Rhinatrema bivittatum* | Gymnophiona | LWS | XM_029609805.1 | NA |
|  |  | RH1 | XP_029453475.1 | NA |
| *Rhinella arenarum* | Anura | LWS | TSA: GHCG01033884 | NA |
|  |  | RH1 | TSA: GHCG01029964 | NA |
| *Rhinella marina* | Anura | LWS | TSA: GFMT01011396 | NA |
|  |  | RHO | TSA: ONZH01010496 | NA |
| *Salamandra infraimmaculata* | Caudata | LWS | SRX2775495 | NA |
| *Salamandra salamandra* | Caudata | LWS | TSA: GIKK01019808 | NA |
|  |  | RHO | SRX2775496 | NA |
| *Scaphiopus couchii* | Anura | LWS | TSA: GHBO01054696 | NA |
|  |  | RHO | TSA: GHBO01076204 | NA |
| *Scolecomorphus vittatus* | Gymnophiona | LWS | PQ541073 | CAS:Herp:168812 |
| *Spea bombifrons* | Anura | LWS | XM053473919 | NA |
|  |  | RHO | XM053469773 | NA |
| *Spea multiplicata* | Anura | LWS | TSA: GKIA01262822 | NA |
|  |  | RHO | TSA: GKIA01095436 | NA |
| *Staurois parvus* | Anura | LWS | SRX20281871 | NA |
| *Taudactylus pleione* | Anura | LWS | RX20667199 | NA |
| *Typhlonectes natans* | Gymnophiona | LWS | PQ541080 | MVZ:Herp:179731 |
| *Typhlonectes natans* | Gymnophiona | LWS | PQ541081 | MVZ:Herp:179733 |
| *Xenopus laevis* | Anura | LWS | MN820844 | NA |
|  |  | RHO | L07770 | NA |
| *Xenopus tropicalis* | Anura | LWS | NM001102861 | NA |
|  |  | RHO | NM0010973342 | NA |
| *Alligator mississipiensis* | Crocodilia | LWS | XM006269029 | NA |
|  |  | RHO | NM001287282 | NA |
| *Caretta caretta* | Testudines | LWS | XM048832768 | NA |
|  |  | RHO | XM048858495 | NA |
| *Gopherus flavomarginatus* | Testudines | LWS | XM050928050 | NA |
|  |  | RHO | XM050958404 | NA |
| *Mauremys reevesii* | Testudines | LWS | XM039511073 | NA |
|  |  | RHO | XM039547817 | NA |
| *Phyllodactylus unctus* | Squamata | LWS | KU645243 | NA |
| *Podacris muralis* | Testudines | LWS | XM028711359 | NA |
|  |  | RHO | XM028719287 | NA |
| *Pogona vitticeps* | Squamata | LWS | XM020793072 | NA |
|  |  | RHO | XM020813194 | NA |
| *Gallus gallus* | Galliformes | LWS | NM205440 | NA |
|  |  | RHO | NM001030606 | NA |
| *Homo sapiens* | Primates | LWS | NM020061 | NA |
|  |  | RHO | NM000539 | NA |
| *Neoceratodus fosteri* | Ceratodontiformes | LWS | EF526297 | NA |
|  |  | RHO | EF526295 | NA |
| *Protopterus annectens* | Ceratodontiformes | LWS | XM044079282 | NA |
|  |  | RHO | XM044078452 | NA |
| *Amia calva* | Amiiformes | LWS | MN519167 | NA |
|  |  | RHO | MN519151 | NA |
| *Carassius auratus* | Cypriniformes | LWS | L11867 | NA |
|  |  | RHO | XM026250265 | NA |
| *Cyprinus carpio* | Cypriniformes | LWS | AB055656 | NA |
|  |  | RHO | S74449 | NA |

**Supplementary Table 4 – Maximum wavelength absorbance values (λ_max_) for *LWS* and *RH1* in amphibians and other vertebrate species**. Point estimations of λ_max_ for *LWS* and *RH1* were predicted using a machine-learning model (Frazer et al., 2024). The model was trained with the VPOD_vert_het_1.0 dataset, which includes 721 vertebrate opsin genes (13 *UVS*, 167 *SWS*, 8 *MWS*, 83 *LWS*, 237 *RH1*, and 113 *RH2*), incorporating wild-type, mutant, and chimeric opsins with λ_max_ values ranging from 350 to 611 nm. Additionally, we compiled published microspectrophotometry (MSP) data for species included in our machine-learning analysis. In cases where λ_max_ values were unavailable for a particular species, we used data from a closely related species within the same genus, with the species and source explicitly stated. The default chromophore assumed for *λ_max_* estimates is A1; if another chromophore was used, this is noted alongside the reported value. For *Cynops orientalis*, rods exhibited 45% A1 chromophore composition, and absorbance values are reported for both A1 and A2 chromophores.

| **Species** | **λ_max_ *LWS*** | **Average MSP *LWS*** | **λ_max_ *RH1*** | **Average MSP**  ***RH1*** | **Literature** |
| --- | --- | --- | --- | --- | --- |
| *Ambystoma mexicanum* | 556.87 | - | 503.09 | 501.8±1.6 | (Ernst et al., 1978) |
| *Ambystoma tigrinum* | 556.87 | mean 610  565–620^*^ | 503.09 | 506 | (Cornwall et al., 1984; Davies et al., 2012; Jones et al., 1993; Margetts et al., 2024; Perry & McNaughton, 1991) |
| *Bombina bobina* | 558.36 | - | 502.22 | - | - |
| *Bombina pachypus* | 558.36 | - | - | - | - |
| *Bufo bufo* | 554.23 |  | 503.60 | 501.7 | (Govardovskii et al., 2000) |
| *Caecilia orientalis* | 546.87 | - | 493.14 | - | - |
| *Caecilia orientalis_v2* | 508.57 | - | - | - | - |
| *Cynops pyrrhogaster* | 551.47 | 547/601  А1/А 2 | 504.77 | 496 /519 A1/A2 | *C. orientalis* (Korenyak & Govardovskii, 2013) |
| *Geotrypetes seraphini* | 550.58 | - | 492.07 | 486.6±2.2 | (Mohun et al., 2010) |
| *Hyla sarda* | 561.41 | - | 505.10 | 501  503 | *H. regilla*  (Crescitelli, 1959)  *H. cinerea*  (King et al., 1993) |
| *Hynobius chinensis* | 551.83 | - | 504.27 | - | - |
| *Hynobius retardatus* | 544.56 | - | 504.27 | - | - |
| *Ichthyophys bananicus* | 548.48 | - | 503.57 | - | - |
| *Limnodynastes peronii* | 553.68 | - | 500.54 | 502 | *L. ornatus*  *(Partridge et al., 1992)* |
| *Microcaecilia unicolor* | 550.38 | - | 502.77 | - | - |
| *Microhyla fissipes* | 556.27 | - | 500.71 | 504 | *M. olivacea* (Partridge et al., 1992) |
| *Oophaga pumilio* | 545.01 | 561±3 | 499.73 | 491±2 | (Siddiqi et al., 2004) |
| *Pelobates cultripes* | 553.22 | - | 504.50 | - | - |
| *Pyxicephalus adspersus* | 554.27 | - | 501.32 | - | - |
| *Quasipaa spinosa* | 549.46 | - | - | - | - |
| *Rhinatrema bivittatum* | 547.21 | - | 492.13 | 487.5±1 | (Mohun et al., 2010) |
| *Rhinella arenarum* | 551.80 | - | 505.95 | - | - |
| *Rhinella marina* | 551.80 | - | 504.49 | 503.3 | (Govardovskii et al., 2000) |
| *Salamandra infraimmaculata* | 552.91 | - | - | - | - |
| *Salamandra salamandra* | 553.07 | - | 505.29 | - | - |
| *Scaphiopus couchii* | 554.51 | 552 | 513.54 | 512 | (Schott et al., 2024) |
| *Spea bombifrons* | 557.49 | - | 506.97 | - | - |
| *Spea multiplicata* | 554.15 | 556 | 506.97 | 512 | (Schott et al., 2024) |
| *Staurois parvus* | 541.84 | - | - | - | - |
| *Taudactylus pleione* | 557.26 | - | - | - | - |
| *Xenopus laevis* | 552.15 | - | 502.87 | 523/A2 | (Govardovskii et al., 2000) |
| *Xenopus tropicalis* | 553.43 | - | 501.49 | - | - |
| *Alligator mississippiensis* | 542.41 | 566.2±3.8 | 500.74 | 501.2±1.1 | (Sillman et al., 1991) |
| *Gopherus flavomarginatus* | 563.31 | - | 497.45 | - | - |
| *Mauremys reevesii* | 559.48 | 620 | 499.65 | *-* | *M. caspica*  *(Perlman et al., 1994)* |
| *Phyllodactylus unctus* | 539.44 | - | - | - | - |
| *Podarcis muralis* | 561.51 | 562±17 | 498.57 | - | (Martin et al., 2015) |
| *Pogona vitticeps* | 560.12 | - | 493.72 | - | - |
| *Gallus gallus* | 565.30 | 569 | 502.26 | 506 | (Bowmaker & Knowles, 1977) |
| *Homo sapiens* | 554.28 | 560 | 501.18 | 494 | (Morrow et al., 2017) |
| *Neoceratodus fosteri* | 555.83 | 561 | 506.25 | 510 | (Davies et al., 2012) |
| *Protopterus annectens* | 531.39 | - | 500.38 | - | - |
| *Amia calva* | 556.29 | 624±6/Chromophore not stated | 500.69 | 527±2/Chromophore not stated | (Burkhardt et al., 1983) |
| *Carassius auratus* | 556.80 | 625±5/A2 | 498.68 | 516/A2 | (Govardovskii et al., 2000; Hárosi & MacNichol, 1974; Palacios et al., 1998) |
| *Cyprinus carpio* | 550.10 | 619/A2 | 499.08 | 524/A2 | (Govardovskii et al., 2000) |

* A range of 30–40 nm has been reported in spectral variation in red cones in *Ambystoma tigrinum* (Makino & Dodd, 1996).

**Data 1 - *LWS* and *RH1* gene trees.** Matrices, log files, and gene trees for *LWS* and *RH1* sequences, including sequences of representative amphibian species (Anura = 19 spp, Caudata = 8 spp) and 13 caecilian species for *LWS* and 5 for *RH1*. Data were obtained from publicly available genomic resources, PCR amplification and sequencing, and eye transcriptomic sequencing. The trees were constructed using IQ-TREE2, applying Maximum Likelihood (ML) as the optimality criterion. This documentation is stored in the Dryad Digital Repository (DOI: 10.5061/dryad.h18931zxf).

**Data 2 - Alignment of *LWS* and *RH1* with caecilian and relevant species used to estimate *λ_max_* values for both opsins.** The methods used to retrieve *LWS* for this alignment are detailed in the Methods section of the main manuscript and in Section 2 of this document. Our final dataset included 19 species of Anura and 8 species of Caudata (*Ambystoma mexicanum, Ambystoma tigrinum, Bombina bombina, Bombina pachypus, Bufo bufo, Chioglossa lusitanica, Cynops pyrrhogaster, Hyla sarda, Hynobius chinensis, Hynobius retardatus, Limnodynastes peronii, Microhyla fissipes, Oophaga pumilio, Pelobates cultripes, Pyxicephalus adspersus, Quasipaa spinosa, Rhinella arenarum, Rhinella marina, Salamandra infraimmaculata, Salamandra salamandra, Scaphiopus couchii, Spea bombifrons, Spea multiplicata, Staurois parvus, Taudactylus pleione, Xenopus laevis, Xenopus tropicalis*), six species of reptiles (*Alligator mississippiensis*, *Gopherus flavomarginatus*, *Mauremys reevesii, Phyllodactylus unctus, Podarcis muralis, Pogona vitticeps*), 1 species each of birds (*Gallus gallus*) and of mammals (*Homo sapiens*), and 5 species of fish (*Neoceratodus forsteri*, *Protopterus annectens, Amia calva*, *Carassius auratus*, *Cyprinus carpio*). See Supplementary Material Table 1 for accession numbers. This documentation is stored in the Dryad Digital Repository (DOI: 10.5061/dryad.h18931zxf).

**Data 3 - *LWS* and *RH1* alignments with bovine rhodopsin.** Matrices of *LWS* and *RH1* sequences for representative amphibian species (Anura and Caudata), 13 caecilian species for *LWS*, and 5 for *RH1*, along with the bovine rhodopsin sequence (NP_001014890.1). This documentation is stored in the Dryad Digital Repository (DOI: 10.5061/dryad.h18931zxf).

#### **Data 4 - Alignments, trees, control files, and output files from the selection analysis conducted in PAML and HYPHY.** For *LWS*, both the incomplete and complete gene datasets were analyzed, and results from both approaches are provided. For *RH1*, only the complete gene dataset was used, and all analyses were performed using the full matrix. For PAML, we are including the alignments, unrooted trees, control files, and all output files generated during the analyses. For HYPHY, we are providing the alignment uploaded to DataMonkey, along with the JSON files obtained as part of the results. This documentation is stored in the Dryad Digital Repository (DOI: 10.5061/dryad.h18931zxf).

#### **Data 5 - Photomicrographs of specimens examined for morphological characterization of caecilian eyes.** Species IDs, S-book, field and museum numbers (when available), stain, and magnification details are provided for all examined specimens in both Supplementary Table 2 and the document containing the images. This documentation is stored in the Dryad Digital Repository (DOI: 10.5061/dryad.h18931zxf).

#### **Data 6 - PCR fragments of *LWS* from 11 caecilian species.** The amplified region includes a portion of exon I, the intronic sequence between exon I and II, and a fragment of exon II of the *LWS* gene for 11 caecilian species*.* Three species have two samples: *Dermophis mexicanus*, *Gymnopis multiplicata*, and *Typhlonectes natans.* This documentation is stored in the Dryad Digital Repository (DOI: 10.5061/dryad.h18931zxf).

#### **Data 7 – Alignments of cone and rod phototransduction genes from the genomes of anuran, salamander, and caecilian species including the transcriptome of one caecilian species.** We analyzed three frog species (*Xenopus tropicalis*, *Nanorana parkeri*, and *Pyxicephalus adspersus*), two salamanders (*Ambystoma mexicanum* and *Pleurodeles waltl*), and three caecilians (*Microcaecilia unicolor*, *Rhinatrema bivittatum*, and *Geotrypetes seraphini*). Additionally, we searched for these genes in our newly generated eye transcriptome from *Caecilia orientalis* (Supplementary Material Table 1). In caecilians, we were able to retrieve six genes from the rod phototransduction genes: *GNAT1*, *GNB1*, *GNGT1*, *PDE6A*, *PDE6G*, *CNGA1*, and *CNGB1*, with *PDE6B* being the only gene absent and therefore not present in our dataset. In contrast, cone phototransduction genes display more frequent gene loss and a mosaic pattern of presence and absence. Among these, *GNB3* and *CNGA3* are the only genes consistently present across all caecilian species, while *GNGT2* and *CNGB3* are entirely absent from all available caecilian genomes and transcriptomes. The remaining cone genes (*GNAT2,* *PDE6C*, and *PDE6H*) show variable presence across species. See Section 2.3.4 in the main text for a detailed description of the search methodology, and Section 3.4.4, Figure 4, and Table 2 for results on gene presence/absence in caecilian species.

### **Section 2- METHODS**

***2.1 ANCESTRAL CHARACTER STATE RECONSTRUCTION***

We estimated ancestral character states for anatomical features of the eye using character data mainly from (Wake, 1993) , but also, for a few species, from (Norris & Hughes, 1918), Taylor (1968, 1969), and Maciel and Hoogmoed (2011) (see Supplementary Material Table 3). Wake (1993) used her character set only to estimate a caecilian phylogeny because at the time molecular evidence was still fragmentary.  The reconstructions were performed using stochastic character mapping in *phytools* v2.0 (Revell, 2024a) on a time-tree obtained from the (San Mauro et al., 2014) matrix of mitochondrial genomes. The time-tree was inferred using BEAST v2.7.8 (Bouckaert et al., 2019) under a strict clock and the model of DNA evolution HKY. The time-tree was calibrated using three secondary calibrations obtained from (Hime et al., 2021): (1) the most recent common ancestor (MRCA) of Gymnophiona at 295.0 My, (2) the MRCA of Gymnophiona excluding Rhinatrematidae at 115.9 My, and (3) the MRCA of Gymnophiona excluding Rhinatrematidae and Ichthyophiidae at 84.4 My. Taxon sampling between (Wake, 1993) and our phylogeny largely overlaps. However, if a given species from (Wake, 1993) was not included in the phylogeny, we coded it using the most closely related congeneric species available. For example, in *Scolecomorphus*, (Wake, 1993) reports data for two species, *S. uluguruensis* and *S. kirkii*. We coded *S. vittatus* using *S. kirkii* which is the most closely related species (Zhang & Wake, 2009); *S. uluguruensis* was not included in the analyses. To improve the estimates of the ancestral states, we also included representative species of Batrachia (frogs + salamanders), the sister clade of Gymnophiona, following (San Mauro et al., 2014) taxon sampling. We reconstructed two characters. First, eye musculature, with six discrete states: (0) all muscles lost, (1) two muscles present (rectus superior, rectus inferior), (2) three muscles present (rectus superior, rectus inferior, rectus internus), (3) four muscles present (rectus superior, rectus inferior, rectus internus, rectus externus), (4) six muscles present, with superior oblique attenuate, (5) six muscles present. This character was reconstructed under three models: equal rates, all rates different, and ordered. We preferred reconstructing all muscles combined in a single character, because our objective was to understand the gain or loss of eye complexity rather than mapping individual muscles.  The second character was eye exposure, with three discrete states: (0) eye exposed, (1) eye under skin, (2) eye under bone. In *Oscaecilia ochrocephala*, the eyes are covered by bone only in adults, and some individuals can have bone covering one eye only (MHW, pers. obs.). Similarly, *C. gracilis* eyes are not usually covered by bone, but some individuals can have one or both eyes covered by bone (Maciel & Hoogmoed, 2011). We scored both species as “eye under bone”, because we considered that bone covering the eyes, at any life stage, was indicative of decreased visual function. *Scolecomorphus vittatus* has bony orbits and their eye is attached to the tentacle; it was coded as “eye under bone” following (Wilkinson, 1997) . This character was reconstructed under two models: equal rates and all rates different. In both characters, models were compared using AIC and then fitted to the stochastic mapping function according to their weight (*simmap* function). The data were plotted using *phytools* v2.0 (Revell, 2024b). Information on the character states for each individual included in our analysis is provided in Supplementary Material Table 3.

#### ***2.2 MORPHOLOGY AND SEARCH FOR CONE PHOTORECEPTORS***

***2.2.1 Histology.*** Whole heads of adult specimens fixed in neutral buffered formalin and preserved in 70% ethanol were demineralized in 10% formic acid, embedded in paraffin, sectioned frontally or sagittally, mounted on numbered slides, stained alternately with hematoxylin-eosin, picro-ponceau, and Mallory’s or Heidenhain’s azan or Mallory’s trichrome stains (Humason, 1979), cover slipped, boxed, and stored.

***2.2.3 Search for cones and description of the retina.*** Measurements (inner and outer segment lengths of photoreceptors; inner and outer plexiform layer widths) were taken with a stage micrometer. The number of rows of nuclei in the outer nuclear layer (ONL), inner nuclear layer (INL), and ganglion cell layer (GCL) were counted. Given that the sections used are historical and represent both sagittal and frontal sections, consistency in the condition of specimens at the time of embedding and sectioning cannot be guaranteed, nor was it possible to make measurements from a standardized reference point. Instead, representative images which contained the largest cross section of the eye while showing good preservation of retinal morphology were selected for cell counts and measurements of retinal layers. Measurements are sometimes given in ranges to account for variation in cell size and spatial variation across the retina. Multiple slides flanking the selected sections and multiple specimens of the same species (when available) were examined to ensure consistency in measurements.

***2.3 FURTHER VALIDATION OF A CAECILIAN* LWS *AND ASSESSMENT OF ITS POTENTIAL FUNCTIONALITY***

***2.3.2 PCR of* LWS *in additional caecilian species.*** For the QCAZ samples, DNA was extracted from liver or muscle tissues preserved in 95% ethanol, following standard guanidine thiocyanate extraction protocols. DNA from MVZ and CAS samples was extracted from liver or muscle preserved in 95% ethanol by salt extraction. PCR conditions were as follows: denaturation at 94˚C for 2 min, followed by 38 amplification cycles (94˚C for 30 s, 54˚C for 30 s, 72˚C for 1 min), and a final extension at 72˚C for 7 min. Resultant amplicons were purified with ExoSAP-it (Applied Biosystems; Waltham, Massachusetts).

***2.3.3 Reviewing available genomes*** For NCBI sequence references, we searched annotated genes, transcripts, and mRNAs within the nucleotide collection (nr/nt) that had a unique accession number that matched Gymnophiona-OPN1LW using BLASTn with of e-values of <10⁻^20^ as the threshold, and we restricted our search to vertebrates. The sequences retrieved included nonstandard nomenclature for the OPN1LW gene with a combination of the following terms: “red sensitive opsin”, “opsin 1”, “red sensitive cone opsin”, “red cone visual pigment”, “red cone opsin”, “long wavelength sensitive opsin”, and “LWS opsin”. The list of all species with OPN1LW for our analysis is provided in Supplementary Material Table 1 and includes only representative vertebrates, with an emphasis on amphibians.

For Gymnophiona, we found three sequences: *Microcaecilia unicolor* OPN1LW (XM_030192059.1), *Geotrypetes seraphini* (XM_033922369.1) and *Rhinatrema bivittatum* (XM_029609805.1). All three sequences correspond to an automatic annotation of the *LWS* gene on the basis of the NCBI eukaryotic gene prediction tool Gnomon v8.2 and 8.4 (Souvorov et al., n.d.) as the ‘best-placed RefSeq’. For Dipnoi, we found two sequences, one for N*eoceratodus forsteri* (EF526297.1) for an *LWS* mRNA experiment and another from an *LWS* gene prediction derived from the *Protopterus annectens* genome and annotated as medium wave sensitive (MWS) opsin 1-like (XM_044079282.1). We were unable to find an annotated sequence for OPN1LW in *P. annectens*, yet the MWS (XM_044079282.1) was the closest match to the *LWS* found in *N. forsteri* (e-value <10⁻^150^, similarity of 84.55% and 91% cover), and in bony fish and other vertebrates (including amphibians) with e-values <10⁻^140^, similarity of ~77% and ~88% cover. Therefore, we cannot rule out that this *MWS* (XM_044079282.1) of *P. annectens* is an incorrect annotation or reconstruction for an *LWS*, as it was assigned by the automatic pipeline of Gnomon v9.0 (Souvorov et al., n.d.) as the ‘best-placed RefSeq’ without any further verification. We further investigated the *Protopterus annectens* genome (PAN1.0, GCF_019279795.1) to reconstruct a *LWS* transcript using a BLASTn search with *N. forsteri* (EF526297.1) as a reference against PAN1.0. This approach resulted in a better *LWS* reference for *P. annectens* located on chromosome 8 (NC_056741.1), which was included in our analyses. The most important difference between the XM_044079282.1 *MWS* predicted gene, and the *P. annectens* genome reconstruction is in exon 1, with the latter being more similar to the *LWS* found in *N. forsteri* and in other vertebrates.

For unannotated transcriptomes, we searched *LWS* orthologs using Gymnophiona-OPN1LW as a reference within the transcriptome shotgun assembly (TSA) database and restricted our search to Amphibia and Dipnoi. Unannotated transcripts retrieved for further analysis were those that closely matched Gymnophiona-OPN1LW using BLASTn and BLASTx with <10⁻^20^ e-values as the threshold. The list of TSA accession numbers from amphibians that were homologous to Gymnophiona- OPN1LW is provided in Supplementary Material Table 1. With TSA and NCBI OPN1LW sequence references, we performed an alignment guided by codon translation, which was trivial except for exon 1. This first exon significantly differed in length between *Microcaecilia unicolor* OPN1LW (XM_030192059.1) and the other two Gymnophiona: *Geotrypetes seraphini* (XM_033922369.1) and *Rhinatrema bivittatum* (XM_029609805.1). Most vertebrates have 37-39 residues in OPN1LW exon 1, but only one Gymnophiona (*M. unicolor* OPN1LW) has such an exon 1 in that range. In contrast, *G. seraphini* (XM_033922369.1) was much shorter with 11 residues, as was *Rhinatrema bivittatum* (XM_029609805.1) with 32 residues. Given the relatively conserved length of exon 1 across vertebrates, we reconstructed the *LWS* gene from the published genomes of the three Gymnophiona: *Geotrypetes seraphini* (GeoSer1.1, GCF_902459505.1), *Rhinatrema bivittatum* (aRhiBiv1.1, GCA_901001175.1), and *Ichthyophis bannanicus* (*I. kohtaoensis*) (NWPU_BC_v1, GCA_033557465.1).

With each genome above, we performed a BLASTn against *Microcaecilia unicolor* OPN1LW (XM_030192059.1) as reference. For *Geotrypetes seraphini* (aGeoSer1.1), we were able to find the full sequence *LWS* (including exon 1) in chromosome 1 (NC_047084.1). For *Rhinatrema bivittatum* (aRhiBiv1.1), we were able to find 89% of the *LWS* sequence in the CAAJIF010007075.1 contig, which corresponds to chromosome 1 (NC_042615.1). However, exon 1 was not recovered. We repeated this research against the newer version of *Rhinatrema bivittatum* (aRhiBiv1.2, GCA_901001135.2), the results were again for a match in chromosome 1 (LR584387.1), but no exon 1 was recovered as it was with aRhiBiv1.1. For *Ichthyophis bannanicus* (*I. kohtaoensis,* NWPU_BC_v1), we were able to find the full *LWS* sequence (including exon 1) on chromosome 2 (CM066032.1, JAWIIG010000002.1).

Sequences were identified as *LWS* orthologs if they closely matched the Gymnophiona reference for *Microcaecilia unicolor* *OPN1LW* (XM_030192059.1). Homology was assessed using BLAST+, and sequences were included if they contained all six exons (as in the human *OPN1LW* gene, NM_020061.6) or if most exons had e-values of <10⁻^20^.

#### ***2.4 REASSESSING EVIDENCE FOR* LWS *LOSS: PRIMER ALIGNMENT ANALYSIS***

A previous attempt to amplify *LWS* (also known as *OPN1LW*) from two species of caecilians (*Ichthyophis* cf. *kohtaoensis* and *Typhlonectes natans*) using eye cDNA from adult specimens was unsuccessful (Mohun et al., 2010). Given that previous work reported a putatively functional copy of *LWS* in three other species of caecilians (Lin et al., 2024), we wanted to verify whether a mismatch between primer and template could have resulted in a failed PCR. Thus, we used the 'water' local alignment tool from EMBOSS (Rice et al., 2000), which implements the Smith-Waterman algorithm, to review potential alignments between primers from Mohun et al. (2010) and cDNA sequences of three published caecilian *LWS* genes (*Rhinatrema bivittatum*, XM_029609805.1; *Microcaecilia unicolor,* XM_030192059.1; *Geotrypetes seraphini,* XM_033922369.1). We report values for forward primers aligned to the forward sequence and for reverse primers aligned to the reverse complement of the sequence.

### **Section 3- RESULTS**

***3.2 CAECILIAN RETINAL MORPHOLOGY***

***Epicrionops sp.*** (Figure 1A). Rods cylindrical, outer segments similar thickness to inner segments. Outer nuclear layer (ONL) 2–3 rows tightly packed spherical nuclei. Inner nuclear layer (INL) 3–4 rows loosely packed spherical nuclei, ~40 µm. Outer plexiform layer (OPL) inconspicuous due to loosely packed nuclei in INL. Ganglion cell layer (GCL) 1–2 nuclei deep. Lens ellipsoid. Two cone-like photoreceptors highlighted in Figures 2A and 2B.

***Ichthyophis kohtaoensis*** Retina distorted, compressed so lens contacts retina. Rod outer segments shorter in medial retina, longer near retinal margin. Rod outer segment size variability may be due to section distortion; however, similar pattern seen by Himstedt (1995). Rods cylindrical, outer segment thickness similar to inner segments. ONL 1–2 rows tightly packed ellipsoid nuclei. OPL 2–4 µm thick. INL 2–3 rows tightly packed spherical nuclei. GCL 1–2 rows of nuclei. Lens ellipsoid. Cone-like photoreceptor indicated in Figure 2C.

***Dermophis mexicanus*** Rod outer segments thinner at base than inner segments, tapering slightly distally (may be due to stretching of the rods). ONL 2–3 rows of tightly packed spherical nuclei. INL 3–4 rows loosely packed spherical nuclei; ~62 µm. Boundary between ONL and INL not clearly defined, so OPL inconspicuous. GCL 2–3 nuclei thick. Lens ellipsoid. Cone-like photoreceptor indicated in Figure 2D. Figure 2F shows photoreceptor tendency to be stretched, peel from pigment epithelium (PE), a common preservation artifact in retinal sections.

***Typhlonectes compressicauda*** Retina cracked, compressed/shrunken; maintains general shape. Individual photoreceptor morphology not discernible. External limiting membrane (ELM) to PE ~5 µm. ONL 1–2 rows tightly packed ellipsoid or irregularly shaped nuclei; ~20µm. INL 3–4 rows packed nuclei; 25–28 µm. OPL not clearly defined, < 3 µm where visible. GCL 1–2 rows nuclei. Lens spherical.

***Scolecomorphus kirkii*** Rods columnar; outer segments slightly thinner than inner segments, without narrowing at end. ONL 2–3 rows of packed spherical nuclei. INL 2–3 rows moderately packed spherical nuclei, 20–28 µm (smaller near the retinal margin). GCL one nucleus deep; nuclei not contiguous layer throughout length. Lens amorphous, contiguous to GCL. In Figure 2E, a cone-like photoreceptor is indicated with black arrow; photoreceptor similar to double rod indicated with blue arrow.

***3.3 FURTHER VALIDATION OF CAECILIAN* LWS**

**Supplementary Table 5** - Local alignments between *LWS* primers (Mohun et al., 2010) and published *LWS* sequences from three species of caecilians show substantial mismatch between primer and coding sequences. In contrast, primers designed for this study were more similar to published *LWS* sequences.

| **Primer Pair** | **Alignment** | ***G. seraphini*** | ***M. unicolor*** | ***R. bivittatum*** | **Average** |
| --- | --- | --- | --- | --- | --- |
| LWS_117F/542R  (this study) | % Identity  % Similarity  % Gaps | 95.0/90.0  100.0/90.0  0.0/0.0 | 95.0/100.0  100.0/100.0  0.0/0.0 | 95.0/100.0  100.0/100.0  0.0/0.0 | 95.8  98.3  0.0 |
| AMPHLWF1/R1 | % Identity  % Similarity  % Gaps | 57.1/70.0  64.3/80.0  21.4/0.0 | 54.8/73.7  61.3/83.3  19.4/0.0 | 56.8/80.0  59.5/90.0  29.7/0.0 | 65.4  73.1  11.8 |
| AMPHLWF2/R2 | % Identity  % Similarity  % Gaps | 40.4/72.4  44.2/75.9  50.0/10.3 | 63.3/72.4  70.0/79.3  0.0/0.0 | 66.7/72.4  74.1/86.2  3.7/0.0 | 64.6  71.6  10.7 |
| AMPHLWF3/R3 | % Identity  % Similarity  % Gaps | 71.0/77.8  74.2/85.2  6.5/3.7 | 67.6/74.1  70.6/81.5  14.7/3.7 | 73.3/74.1  76.7/81.5  0.0/3.7 | 73.0  78.3  5.4 |
| AMPHLWF4/R4 | % Identity  % Similarity  % Gaps | 74.1/72.4  81.5/75.9  0.0/0.0 | 74.2/70.0  80.6/76.7  3.2/0.0 | 74.2/72.4  80.6/79.3  3.2/0.0 | 72.9  79.1  1.1 |
| ALLOPSINF1/R1 | % Identity  % Similarity  % Gaps | 53.3/58.1  63.3/64.5  0.0/12.9 | 53.3/64.3  63.3/78.6  0.0/0.0 | 56.7/64.3  66.7/78.6  0.0/0.0 | 58.3  69.2  2.2 |
| ALLOPSINF2/R2 | % Identity  % Similarity  % Gaps | 43.8/69.2  52.1/80.8  39.6/0.0 | 41.7/61.5  50.0/73.1  39.6/0.0 | 52.9/69.2  64.7/80.8  14.7/0.0 | 56.4  66.9  15.7 |

**3.4 *FURTHER VALIDATION OF A CAECILIAN* LWS *AND ASSESSMENT OF ITS POTENTIAL FUNCTIONALITY***

***3.4.4 Assessing phototransduction cascade genes to infer* LWS *expression location.***

In *R. bivittatum*, *PDE6A* is 23 amino acids shorter than in other caecilians, and *GNAT1* contains a premature stop codon at position 167; however, the latter is labeled as low quality in NCBI, suggesting a possible annotation artifact. Among other amphibians, *CNGB1* was not detected in *N. parkeri*, and the *P. adspersus* sequence includes a premature stop codon at position 2363 and is also flagged as low quality; this sequence is also labeled as low quality in NCBI (Supplementary Material XX).

In *M. unicolor,* the *GNB3* sequence exhibits substantial amino acid divergence near the C-terminal region (from approximately amino acid 320 onward), despite high similarity with other amphibians across the remainder of the alignment (Table 3, Supplementary Material XX). For *CNGA3* the transcript of *C. orientalis* is incomplete, missing approximately 90 amino acids compared to other amphibian orthologs (alignment starts at position 109; Supplementary Material XX). The remaining 3 genes of the phototransdcution cascade show different patterns of absence/presence state. For example, *GNAT2* was detected only in the genome of one species (*R. bivittatum*) and *PDE6C* is only absent in *M. unicolor,* but incomplete in *C. orientalis* (see Table 3 for more details)

#### ***3.5 SELECTION ANALYSIS***

#### **File S1 – Summary of selection analyses results for *LWS* and *RH1* using BUSTED, FUBAR, and RELAX as implemented in HYPHY.** BUSTED (Branch-site Unrestricted Statistical Test for Episodic Diversification) results are presented in a single sheet for both *LWS* and *RH1* datasets, including analyses with and without caecilians as the foreground. FUBAR (Fast, Unconstrained Bayesian AppRoximation) and RELAX results are provided in separate sheets, one for *LWS* and one for *RH1*. This file is stored in the Dryad Digital Repository (DOI: 10.5061/dryad.h18931zxf).

#### **File S2 – Summary of selection analyses results for *LWS* and *RH1* using models implemented in PAML.** Results are compiled in separate sheets for each gene. For *LWS*, both complete and incomplete dataset results are included, while *RH1* results are presented for the complete dataset. Additional sheets summarize Random-Site Model and BEB results for both genes. This file is available in the Dryad Digital Repository (DOI: 10.5061/dryad.h18931zxf).

#### ***3.5.1* *Selection analyses with incomplete* LWS *dataset***

For *LWS*, we ran analyses on two types of datasets, a complete gene dataset including the full coding sequence of the gene with smaller sampling for Gymnophiona (368 aa; Anura = 19 spp, Caudata = 8 spp, Gymnophiona = 5 spp) and an incomplete dataset with more species for the Gymnophiona clade but fewer sites (132 aa; Anura = 19 spp; Caudata = 8 spp, Gymnophiona = 13 spp). Here, we describe the results obtained from the incomplete dataset of *LWS*. The analyses for *RH1* were always conducted using the complete gene, and those results are presented alongside the results for the complete *LWS* gene in the main text.

Consistent with the complete *LWS* gene analysis, we found that the M3 model provided a better fit than the M0 model (likelihood ratio [LR] = 347.076, P < 0.0001; File S2), indicating ω rate variation across sites. To test for positively selected sites, we compared the fit of the M8 model (which allows for sites with ω > 1) to the M7 and M8a models. The M8 model was a better fit than M7 (LR = 11.882, P = 0.0026; File S2) but not M8a (LR = 0.797, P = 0.3719; File S2), suggesting a weak signal of positive selection. However, no sites were inferred to be under positive selection with a BEB posterior probability >90%, while one site (position 5; bovine *RH1* numbering) fell below the significance threshold.

Consistent with the complete dataset analysis results, BUSTED found no significant evidence of gene-wide episodic diversifying selection (P = 0.1589, File S1). However, it identified a site (position 154) with evidence ratios (ER) ≥ 100, differing from the site identified by CODEML (position 5) and those reported in the complete dataset analysis (positions 107 and 213) (File S1).

The FEL and FUBAR analyses were used to detect signals of positively selected sites across the entire phylogeny. However, FUBAR did not identify any sites in the incomplete *LWS* dataset when applying a posterior probability threshold of 0.9 (File S1). This result was further confirmed by FEL, which reported no sites under diversifying positive selection, regardless of whether the caecilian clade was used as the foreground or the analysis spanned the entire phylogeny.

Our analysis using RELAX revealed a trend toward relaxation rather than intensification of selective constraint in *LWS* when the crown Gymnophiona and its stem branch were designated as the foreground, though the result was not statistically significant (K = 0.59, P = 0.100, LR = 2.70; File S1). This signal of relaxation was supported by clade model C but not clade model D in the PAML analyses. Specifically, clade model C identified a significant shift in selection pressure for caecilians compared to other amphibians (CmC vs. M2a_rel: LRT = 14.008, P = 0.0002; File S2). However, this was not corroborated by clade model D (CmD vs. M3: LRT = 2.407, P = 0.1208; File S2), suggesting reduced statistical power compared to analyses conducted with the complete gene dataset.

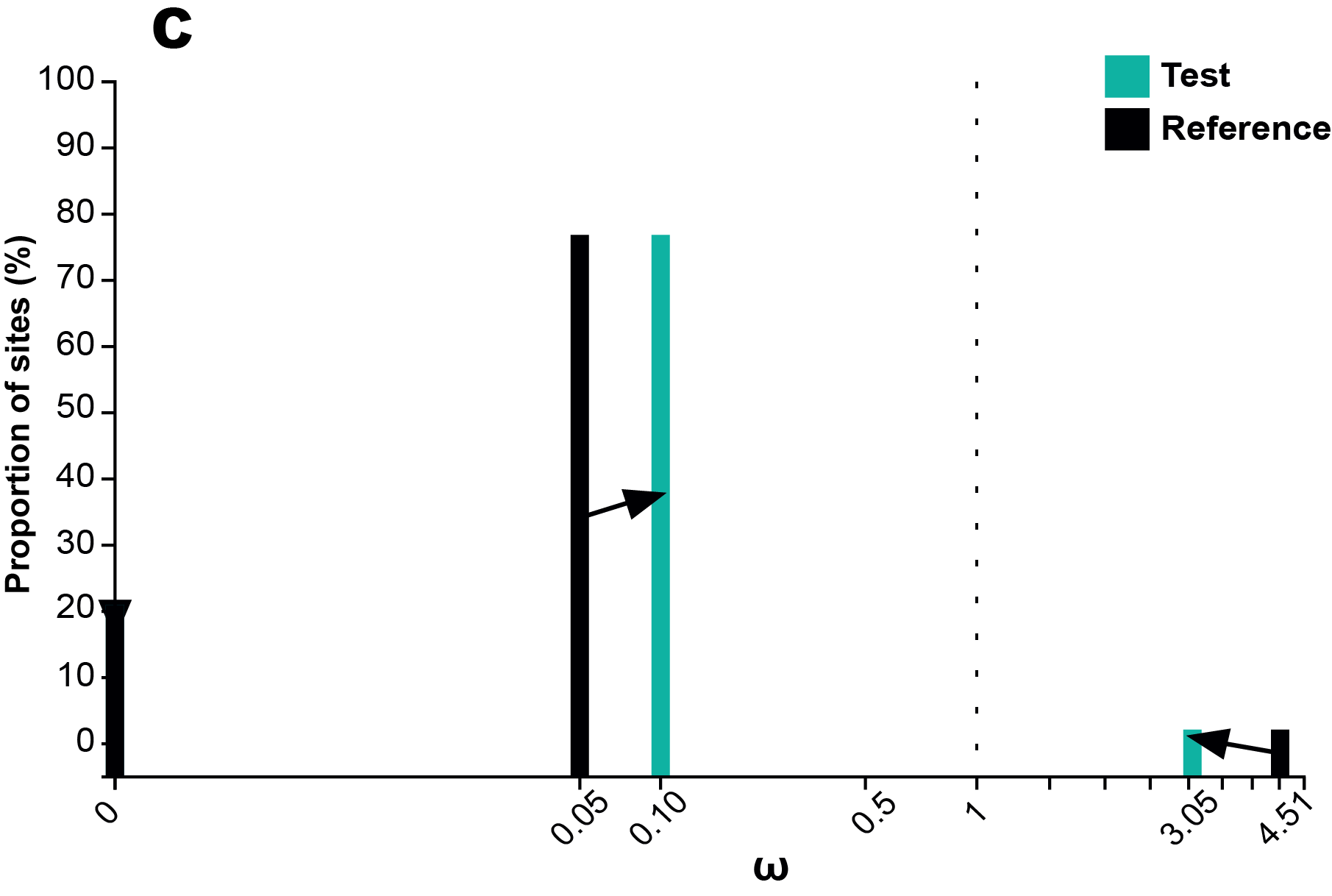

**Supplementary Figure 1.** Results from the RELAX selection analysis indicate that *LWS* in caecilians is undergoing a shift in selective pressure compared to other amphibians (see Supplementary Material File S1). The estimated ω (omega) values for the test group (Gymnophiona, shown in turquoise) compared to the reference group (Anura + Caudata, shown in black) in both ω classes (ω < 1 and ω > 1) are trending toward 1, as indicated by the black arrows. This analysis was performed using the complete dataset (368 amino acids; Anura = 19 species, Caudata = 8 species, Gymnophiona = 5 species).

### **Section 4- DISCUSSION**

***4.2 FUNCTIONAL IMPLICATIONS AND ECOLOGICAL PERSISTENCE OF* LWS *IN CAECILIANS***

#### **Supplementary Table 6 – Amino acid composition of positively selected sites identified in our selection analysis and known spectral tuning sites in *LWS* and *RH1* sequences for Gymnophiona.** Sites are reported using bovine *RH1* numbering. Additionally, we describe their location within the protein’s tertiary structure and report known spectral variants for each site.

| **Site (RH1 numbering)** | **Location** | **Known from** | **Known spectral variants** | **Gymnophiona opsin variants** | |
| --- | --- | --- | --- | --- | --- |
|  |  |  |  | ***RH1*** | ***LWS*** |
| 49 | TMI | Selection analysis | - | - | - |
| 75 | CLI | Selection analysis | - | I | I |
| 83 | TMII | RH1/2 | D/N | N | D |
| 87 | TMII | RH1, SWS1 | V/D/C/A | V | T |
| 90 | TMII | RH1, SWS1 | G/S/C | G | A |
| 94 | TMII | RH1/SWS2 | A/S/C | T | S |
| 96 | TMII | RH1 | Y/V | Y | I/V |
| 100 | ECI | LWS | S/Y | N | F |
| 102 | ECI | RH1 | Y/F | Y | Y/F |
| 107 | ECI | Selection analysis | - |  |  |
| 113 | TMIII | RH1, SWS1 | E/D | E | E |
| 118 | TMIII | RH1, SWS1/2 | S/T/A/G | T | S |
| 119 | TMIII | RH1 | L/F | L/I | V |
| 122 | TMIII | RH1, SWS2 | E/I/Q/M | E | I |
| 124 | TMIII | RH1 | A/S/G/V | A | G/A |
| 132 | CLII | RH1 | A/S/V | A | A/S |
| 164 | TMIV | RH1, RH2, LWS | S/A | A | S |
| 181 | ECII | LWS | H/Y | E | H |
| 194 | ECII | RH1 | P/R | L | G |
| 195 | ECII | RH1 | N/A | K | S/N |
| 207 | TMV | RH1, RH2/SWS2 | M/L | M | L |
| 208 | TMV | RH1 | F/Y | F | M |
| 211 | TMV | RH1 | H/C | H | C |
| 214 | TMV | LWS | T/I | I | I |
| 217 | TMV | LWS | S/A/G/I | L/S/T | S/T |
| 261 | TMVI | RH1, SWS2, LWS | F/Y | F | Y |
| 265 | TMVI | RH1, SWS1/2 | W/Y | W | W |
| 269 | TMVI | SWS2, LWS | A/S/T | A | T |
| 292 | TMVII | RH1/2, LWS, SWS2 | A/S | A/S | A |
| 293 | TMVII | LWS | Y/F | F | Y |
| 295 | TMVII | RH1 | A/S | A | A |
| 299 | TMVII | RH1 | A/S | A | A |
| 300 | TMVII | RH1 | I/T/L | I | T |
| 317 | C-T | RH1 | A/I/M/T | M | I |

#### ***4.3 CAECILIAN EYE MORPHOLOGY***

The socket covered by bone, as well as skin, muscle, and connective tissue, may be a labile character for some taxa, at least the Caeciliidae. *Oscaecilia ochrocephala* have been found with only one of the two sockets covered by bone (MHW, pers. obs.). Furthermore, the bone-covered socket has been observed in some populations of *Caecilia gracilis*, but not in many others. This then poses an evolutionary developmental and a taxonomic conundrum. Wake (unpub.) has conjectured that the bony extension of the maxilla over the socket is a late-developing feature (based on developmental series), so some adults may simply have incomplete peramorphosis of the character. The situation for *C. gracilis* poses a different problem: the genus *Oscaecilia* was described by (Taylor, 1968) based on the single character of the covered socket to distinguish it from its sister genus, *Caecilia*. Consequently, should *C. gracilis* be split, with the populations with covered sockets designated a separate species and assigned to *Oscaecilia*? Or should *Oscaecilia* be sunk, because of the apparently labile nature of the character? Skull development and adult osteology have been examined for very few species of either genus, so knowledge of the presence/absence of the character is largely assumed based on visibility of the eye to the observer (*Caeciia*) or its invisibility (*Oscaecilia*). A study based on comparative morphology and molecular genetics of a large species sample size is necessary to solve this conundrum.

Similarly, all Scolecomorphidae appear to have bone-covered sockets, and, in *Scolecomorphus,* the species studied have the eye not in the socket but embedded in the tentacle. This presumably occurred before the socket covering was completed, and as the tentacle developed, co-opting some eye structures (Billo & Wake, 1987). But no evidence has been presented that species of *Crotaphatrema*, the other genus in the family Scolecomorphidae, have their eyes embedded in the tentacle. Species descriptions, etc., state only that the eye is “not visible externally.” So it is not yet known whether the *Crotaphatrema* condition is like that of *Scolecomorphus*, or that of other caecilians with the orbit in the socket that is covered by the bone.
